# Cells Prioritize the Regulation of Cell Mass Density

**DOI:** 10.1101/2024.12.10.627803

**Authors:** Jinyu Fu, Qin Ni, Yufei Wu, Anoushka Gupta, Zhuoxu Ge, Hongru Yang, Yasin Afrida, Ishan Barman, Sean X. Sun

## Abstract

A cell’s global physical state is characterized by its volume and dry mass. The ratio of cell mass to volume is the cell mass density (CMD), which is also a measure of macromolecular crowding and concentrations of all proteins. Using the Fluorescence eXclusion method (FXm) and Quantitative Phase Microscopy (QPM), we investigate CMD dynamics after exposure to sudden media osmolarity change. We find that while the cell volume and mass exhibit complex behavior after osmotic shock, CMD follows a straightforward monotonic recovery in 48 hours. The recovery is cell-cycle independent and relies on a coordinated adjustment of protein synthesis and volume growth rates. Surprisingly, we find that the protein synthesis rate decreases when CMD increases. This result is explained by CMD-dependent nucleoplasm-cytoplasm transport, which serves as negative regulatory feedback on CMD. The Na^+^/H^+^ exchanger NHE plays a role in regulating CMD by affecting both protein synthesis and volume change. Taken together, we reveal that cells possess a robust control system that actively regulates CMD during environmental change.

## Introduction

How mammalian cells coordinate growth and maintain size across changing environments is a fundamental question in cell biology.^1,2^ Recent advances in single-cell methods have enabled precise measurements of cell volume, mass, and cell mass density (CMD). For instance, microfluidic-based methods such as the Fluorescence eXclusion method (FXm) can accurately measure cell volume in the 10-100 femtoliter range.^3–7^ Micro-cantilever methods have been employed to measure cell dry mass in the femtogram (fg) range.^8–10^ Optical methods such as Quantitative Phase Microscopy (QPM) can also accurately determine the dry mass (without water) of single cells with picogram sensitivity.^11–13^ Therefore, an unprecedented window is opening for examining live cell mass and volume, and also the cell mass density (mass/volume). In this paper, utilizing a combination of FXm and QPM, we quantitatively explore CMD regulation after media osmolarity change. We discover that while complex changes occur in cell volume and mass after osmotic shock, cells prioritize CMD recovery in a straightforward manner, suggesting that there is a simple control algorithm for CMD.

The composition of the mammalian cell cytoplasm has been explored and reviewed.^14–18^ It is known that ions contribute the most to the cytoplasmic osmolarity, while proteins (ribosomes and non-ribosomal proteins) are the bulk of cells’ mass. Recent studies have explored many aspects of cell volume and cell mass dynamics in growing cells.^3,6,7,14,18–21^ Accurate and sensitive measurements of cell mass and volume change in different osmotic and mechanical conditions have been performed.^7,22–26^ Mechanisms of sensing cell volume change during osmotic shock have been proposed.^2,7,27–31^ Cell volume and mass regulation have been shown to profoundly impact cell growth and dynamics. For instance, it has been proposed that mammalian cells divide using a G1/S size checkpoint.^1,32^ Cells can quickly alter their volume and swell during mitosis^21,33^ and neutrophil activation.^34^ Moreover, the total protein concentration itself is an important parameter that influences all biochemical processes by affecting intracellular diffusion, transport, and phase separation.^18,31,35,36^ Thus, the ratio between cell mass and volume, or CMD, is also expected to be a critical variable. CMD should be regulated in a narrow range, and volume and mass regulation should be coupled to the CMD to achieve consistent biological function. Indeed, CMD seems to be cell cycle-independent, and small variations in CMD during cell growth have been observed.^37^ Alterations in CMD have been observed in aneuploid cancer cells^38^, and may be a cause of cellular senescence.^39–41^

In this paper, we also find that while the cell mass and volume can vary significantly across different tissue cell types, the CMD is relatively uniform, except for a few cancer cell types. When cells are exposed to hypertonic shocks, CMD is quickly disrupted as the cell volume initially decreases due to rapid water loss. However, over a longer time scale (∼10 hours), the cell volume and mass begin to increase at an unequal rate. There is more volume increase than mass increase, resulting in a regulatory decrease in the CMD for a 48-hour period. We discover that this active regulation of CMD is not due to cell cycle redistribution. Rather, all cells uniformly regulate their CMD such that the CMD of dividing cells follows the same trend. The Na^+^/H^+^ exchanger (NHE), whose activity causes cell swelling,^42,43^ is involved in regulating CMD, together with the NHE-actin linker ezrin.^7,44,45^ Moreover, protein synthesis is downregulated during CMD recovery, and the protein synthesis rate depends on the CMD. We observe that, counterintuitively, higher CMD results in a lower protein synthesis rate. This result can be explained by changes in the nucleoplasm to cytoplasm (N-C) transport, measured using fluorescence recovery after photobleaching (FRAP). Changes in N-C transport decrease the cytoplasmic ribosome number and the protein synthesis rate. Taken together, we find that there is a system of cell components that actively regulates CMD. This system involves the actin cytoskeleton, NHE, NKCC, and their associated signaling pathways. The interactions that generate this active regulation may be complex, but the net effect on CMD is surprisingly simple. The system behaves similarly to a proportional-integral-differential (PID) controller, with proportional error as the most dominant signal. This system adjusts CMD by simultaneously controlling cell water content (through the action of ion exchangers) and protein synthesis. Our work shows that CMD regulation plays an important role in cell adaptation to environmental changes. Regulators of CMD constitute a major cell system that affects protein concentration in the nucleoplasm and the cytoplasm, governs protein synthesis and regulates cell growth.

## Results

### Cell mass density (CMD) is tightly regulated across cell types and the cell cycle

To obtain quantitative CMD measurements, we employed FXm (Fig. 1A, C) and QPM (Fig. 1B,D). FXm uses a microfluidic channel infused with high molecular weight dextran dye that is impermeable to the cell membrane. Since cells exclude the dye from their cytoplasm, the cell volume can be measured from an epifluorescence image. Therefore, FXm accurately measures the physical volume (𝑉) of the cell.^46^ QPM measures the change in the refractive index of the cytoplasm with respect to the media, and since the refractive index is proportional to the solute mass density (Fig. s1D), the total dry mass (𝑀) and CMD can be calculated from the cell volume and the optical volume (Materials and Methods). ^11–13^ Both methods are high throughput and can generate large data sets in a single experiment. In practice, an FXm measurement yields the population volume average, 〈𝑉〉, and a separate QPM measurement yields the population mass average, 〈𝑀〉. The cell population CMD is then 𝜌 = 〈𝑀〉/〈𝑉〉 (Fig. s1A). To obtain statistically significant results, more than 74 cells were measured in each technical repeat (single data points in figures), and 2 technical repeats were performed for each biological repeat for each cell type.

**Figure 1:**
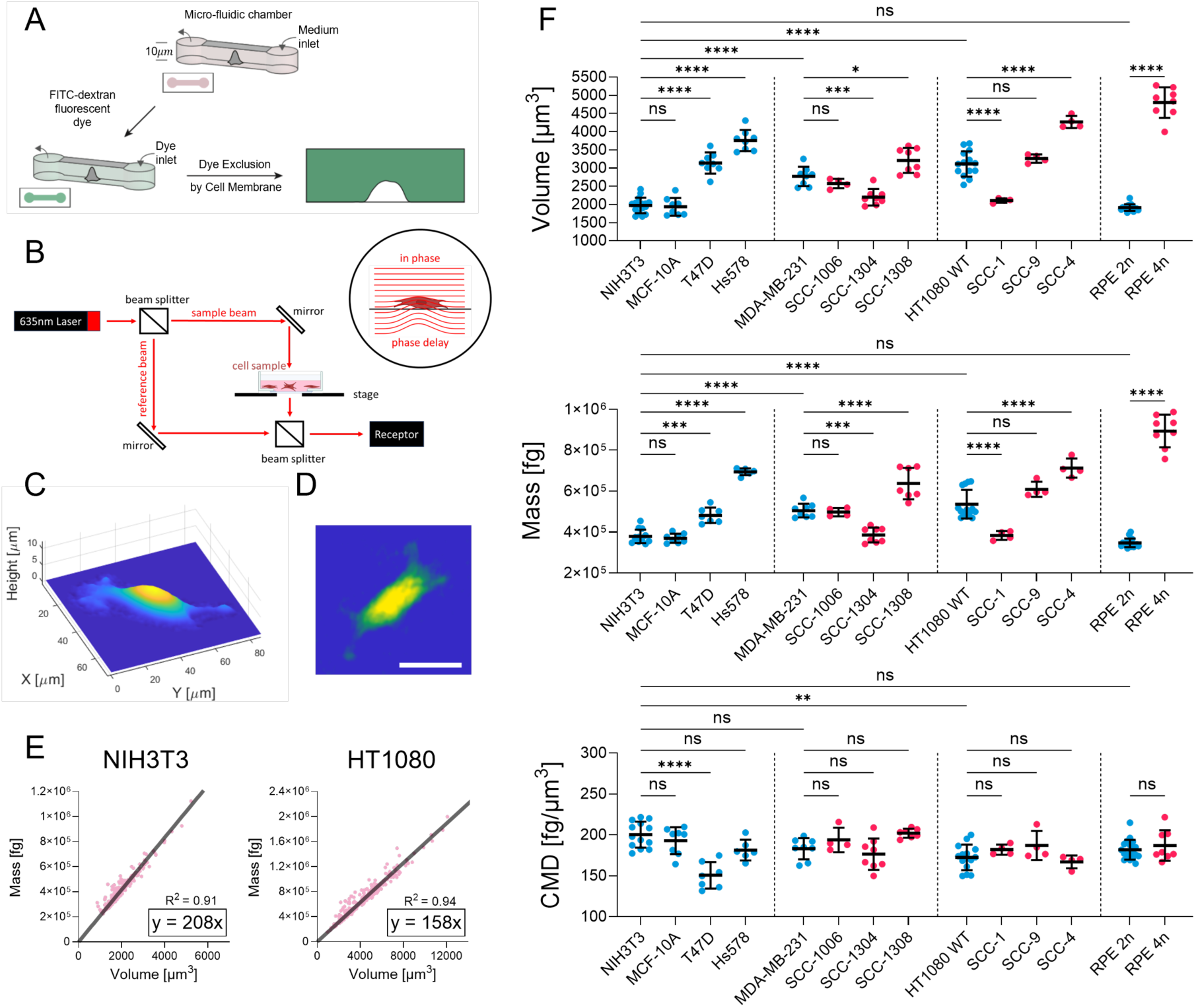
Cell mass density is tightly regulated. (A) A schematic of the Fluorescence Exclusion method (FXm). (B) A schematic of the Quantitative phase microscopy diagram (QPM). (C) A representative FXm 3D cell shape reconstruction. (D) A representative QPM cell image. Pixel color represents the relative optical path difference. (E) Single-cell mass versus volume for NIH3T3 and HT1080 cells from 3D tomographic microscope measurements. Each pink dot represents a single cell. n ≥ 262 cells for each condition. Dark gray trendlines passing through the origin are linear fittings to the data points, which have a slope that represents cell densities. (F) Cell volume, mass, and density for different cell types, single-cell clones (SCCs), and tetraploids (4n) measured by FXm and QPM. Each dot represents the populational average results of one technical repeat. Wild-type cells are presented in blue, and single-cell clones and tetraploids are represented in red. N ≥ 4 repeats for each condition. (F) Data are presented as the mean ± standard deviation. One-way ANOVA tests followed by Dunn-Šidák’s multiple-comparison test were conducted between data sets. *p^ns^* > 0.05, *p*\* ≤ 0.05, *p*\*\* ≤ 0.01, *p*\*\*\* ≤ 0.01, *p*\*\*\*\* ≤ 0.0001. (D) Scale bars, 30 μm.

To verify our mass density measurements, we also employed a 3D optical density tomographic (ODT) microscope to acquire single cell 3D phase images and quantified single cell volume, mass, and density (Fig. s1B,E).^47^ Plotting single cell volume versus single cell mass shows that volume-mass scaling is linear across all cell sizes (Fig. 1E). Since there is a mixture of G1 and G2 cells in the experiment, this data indicates that there is no significant density variation across different cell cycle stages, in agreement with previously reported results.^37^ CMD is the slope of the linear mass-volume line, which gives 208 fg/µm³ for NIH-3T3 cells, in quantitative agreement with our high-throughput population measurement (Fig. 1F).

We then extended our measurements to include multiple cell types, comparing human and mouse cells, normal and cancer cells, as well as cells with different ploidies (Fig. 1F). To obtain large scale data, we utilized the population volume, mass, and density, combining FXm with QPM. A screening of different cell types revealed that while cell volume and mass can vary significantly across different cell types, CMDs were consistently around 180-200 fg/µm³. For example, the tetraploid hTERT-immortalized retinal pigment epithelial cells (RPE-1) had twice the volume and mass of the diploid cells, but their mass densities were the same. These results suggest that CMDs across different cell types show less variation than cell volume or cell mass. An exception was observed in T47D human breast infiltrating ductal carcinoma cells, which had a lower CMD at 150 fg/µm³, suggesting a relatively swollen cytoplasm.

We also analyzed the volume and density of single-cell clones isolated from HT1080 containing different ploidy numbers (Fig. s1F) and single-cell clones isolated from MDA-MB-231 cells having different metastatic potentials.^48^ We again found that while cell sizes of single-cell clones could differ significantly from their parental cells, the same CMD was generally maintained. Note that it has been shown that the single cell clones do not maintain the same ploidy after extended culture.^48^ CMD was also consistent between diploid and tetraploid RPE-1 cells. Together, these results indicate that most cell types preserve a similar and relatively stable mass density independent of the cell cycle stage or ploidy.

### Hypertonic shock disrupts cell mass, volume, and CMD

To understand how cell volume and mass are coupled to maintain a stable cell mass density, we applied hypertonic shock to disrupt cell volume and monitored changes in mass and density. Previous work showed that osmotic shock is an excellent way to disrupt cell volume and CMD for an extended period.^7,37,49,50^ By adding membrane impermeable osmolyte sorbitol, we increased the media osmolarity by 45% and monitored NIH-3T3 volume, mass, and CMD changes over several days (Fig. 2A,B, Fig. s1C). Note that all nutrients and FBS concentrations were maintained the same as the isotonic control. At short timescales (<1hr), the sudden increase in extracellular osmolarity caused water efflux and an immediate 25% decrease in cell volume. However, large intracellular molecules, such as proteins, amino acids, and metabolites that cannot freely cross the plasma membrane remained, giving a constant cell mass (Fig. 2C). Consequently, CMD increased by 25% immediately following the hypertonic shock. We observed no significant recovery in volume or density within 1 hour after hypertonic shock. After that, cell volume started to recover, reaching its isotonic volume at around 7 hours post-shock. On the other hand, cell mass continuously increased during this period. These led to a CMD recovery to ∼10% higher than its isotonic value.

**Figure 2:**
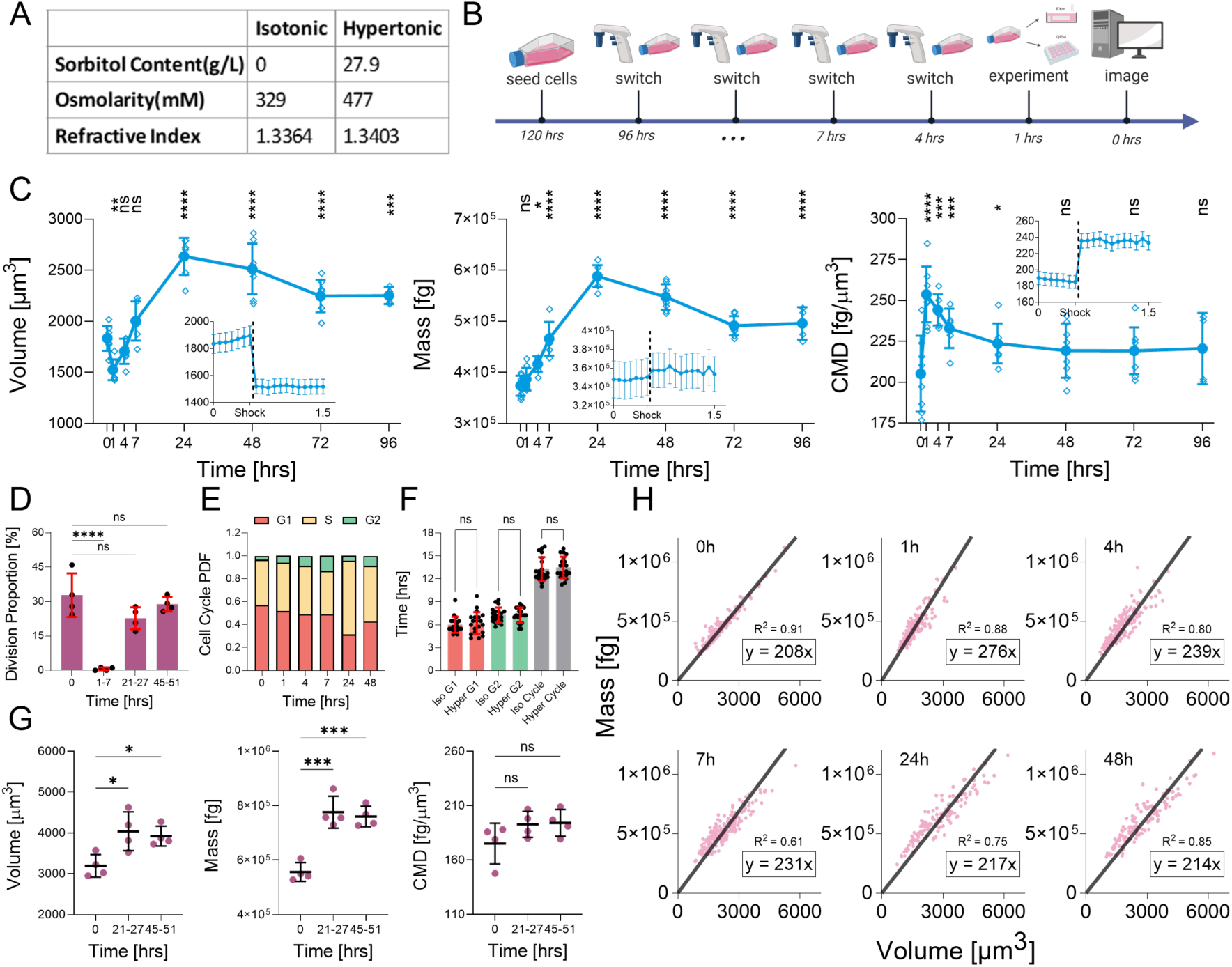
Hyperosmotic shock triggers mass density changes and cell cycle-independent CMD recovery. (A) Table showing properties of isotonic and 145% hypertonic culture medium. (B) A schematic of the hypertonic shock experiment time course. (C) NIH-3T3 cell volume, mass, and density over time before hypotonic shock (0 hrs) and during hypertonic shock (1-96 hrs). Each small circle represents one repeat (N ≥ 4). Inset in (C) are imaged with a 5-minute frame rate, from 30 minutes before to 60 minutes after the hypertonic shock (n ≥ 151). The dashed line indicates the time point when hypertonic shock is introduced. (D) Fraction of cells that divided during a 6-hour time window before and after exposure to hypertonic shock (N = 4). (E) Cell cycle distribution before and after exposure to hypertonic conditions determined from 3T3-FUCCI cells. (F) G1, G2, and the total cell cycle lengths of the first-generation daughter cells born after hypertonic shock (n = 20). (G) Volume, mass, and density of dividing cells in the isotonic environment, and at 24 and 48 hrs in hypertonic condition (N = 4). CMD of dividing cells follows the same trend as in (C). (H) Single-cell mass versus volume before and after exposure to hypertonic shock (n ≥ 202). Trendlines are fits to the data. Slopes represent CMDs. (C, D, F, G) Data are presented as the mean ± standard deviation. One-way ANOVA tests followed by Dunn-Šidák’s multiple-comparison test were conducted between data sets. *p^ns^* > 0.05, *p*\* ≤ 0.05, *p*\*\* ≤ 0.01, *p*\*\*\* ≤ 0.01, *p*\*\*\*\* ≤ 0.0001.

At 96 hours post-hypertonic shock (Fig. 2C), however, we observed that both cell volume and mass continuously increased to values much larger than those in an isotonic environment. Both cell volume and mass peaked at 24 hours post-shock and remained 23% above the initial size after 72 hours. In contrast, CMD, which initially increased, continuously decreased after the shock, eventually stabilizing at a steady state that is similar to the isotonic control after 48 hours.

Together, these observations reveal that volume, mass, and CMD do not follow the same recovery dynamics. Importantly, the monotonic recovery of mass density, as opposed to the non-monotonic recovery of volume and mass, suggests that cells prioritize CMD regulation.

### Density recovery post hypertonic shock is cell cycle independent

After observing the persistent mass and volume increase and density recovery after hypertonic shock, we asked whether these changes were the results of cell cycle redistribution. We first quantified the fraction of cells that divided in a 6-hour time window before and after hypertonic shock (Fig. 2D). We found that almost no cells divided between 1 to 7 hours post shock, in contrast to ∼30% dividing in isotonic media, suggesting that hypertonic shock delayed cell cycle progression to G2/M.^51^ This accumulation of relatively large cells in the late G2 phase explains the population average cell volume and mass increase between 1 to 7 hours in hypertonic environment. Interestingly, the cell division rate recovered after 24 and 48 hours. We also measured the volume, mass, and CMD of dividing cells 10 minutes before cell separation at 24- and 48-hours post-shock (Fig. 2G, Fig. s1G). We found that the volume, mass, and CMD of these mitotic cells increased in a similar way as interphase cells. The results in mitotic cells again indicated that hypertonic shock-induced CMD change is independent of the cell cycle. Note that the measurement was taken during mitotic swelling, therefore the mass density is slightly lower than the population average. These observations also showed that the hypertonic shock-induced G2/M cell cycle arrest is reversible, and very few cells undergo quiescence or senescence. Examining cell cycle distribution using Propidium Iodide DNA content staining and 3T3- FUCCI cells further supported this observation (Fig. 2E,F).^52–54^ These results together show that while hypertonic shock affects the cell cycle, especially at 1-7 hours post-shock, the density recovery is independent of cell cycle change.

Finally, to further confirm our results, we again used 3D ODT microscopy to image single-cell size and density in isotonic conditions and at 1, 4, 7, 24, and 48 hours in the hypertonic condition (Fig. 2H). The cell size and density measurements from single-cell analysis were consistent with our population-level results, and no cell cycle dependence was observed. These results confirmed that cells of different sizes recover their CMD at a similar rate, and the recovery is independent of the cell cycle.

### NHE increases cell volume growth rate and plays a role in regulatory CMD decrease

Since CMD is the ratio between cell dry mass and volume, after a sudden density increase, cells can both increase the volume growth rate and decrease the mass growth rate to recover density. This simultaneous control is exactly what we observed in Fig. 2C. To check for cell components responsible for volume change, we examined two major ion exchangers, the sodium-hydrogen antiporter (NHE) and the Na-K-Cl cotransporter (NKCC), whose activity can both increase intracellular osmolarity and cell volume. NHE, when activated, transports one sodium into the cytoplasm and one proton out. CO₂ freely diffuses across the cell membrane, and when a proton is exported, it is stoichiometrically converted into a bicarbonate ion, thereby increasing cytoplasmic osmolarity. Thus, NHE activation increases the cytoplasmic pH and swells the cell.^7,42,43^ Moreover, the MEK/ERK pathway is critical for growth, proliferation, and signal transduction that interacts with NHE.^55^ Ezrin, a cytoskeletal protein that activates NHE,^7,44,45^ is also overexpressed in hypertonic conditions.^56^ NKCC transports one sodium, one potassium, and two chloride ions into the cell, and increases the cell volume.^43^ Thus, it is likely that NHE and NKCC will play a role in the observed cell volume dynamics and density recovery.

We first measured the cytoplasmic pH as a proxy for NHE activity using pHrodo at isotonic and 1, 4, 7, 24, and 48 hours in hypertonic conditions (Fig. 3A,B).^57,58^ The intracellular pH increased immediately after hypertonic shock, peaking at 4 hours, and then slightly decreased but remained higher than the isotonic control at 7, 24, and 48 hours. The pH trend suggested that the NHE channel was hyperactivated in hypertonic conditions, with peak activity occurring 4 hours after the shock.

**Figure 3:**
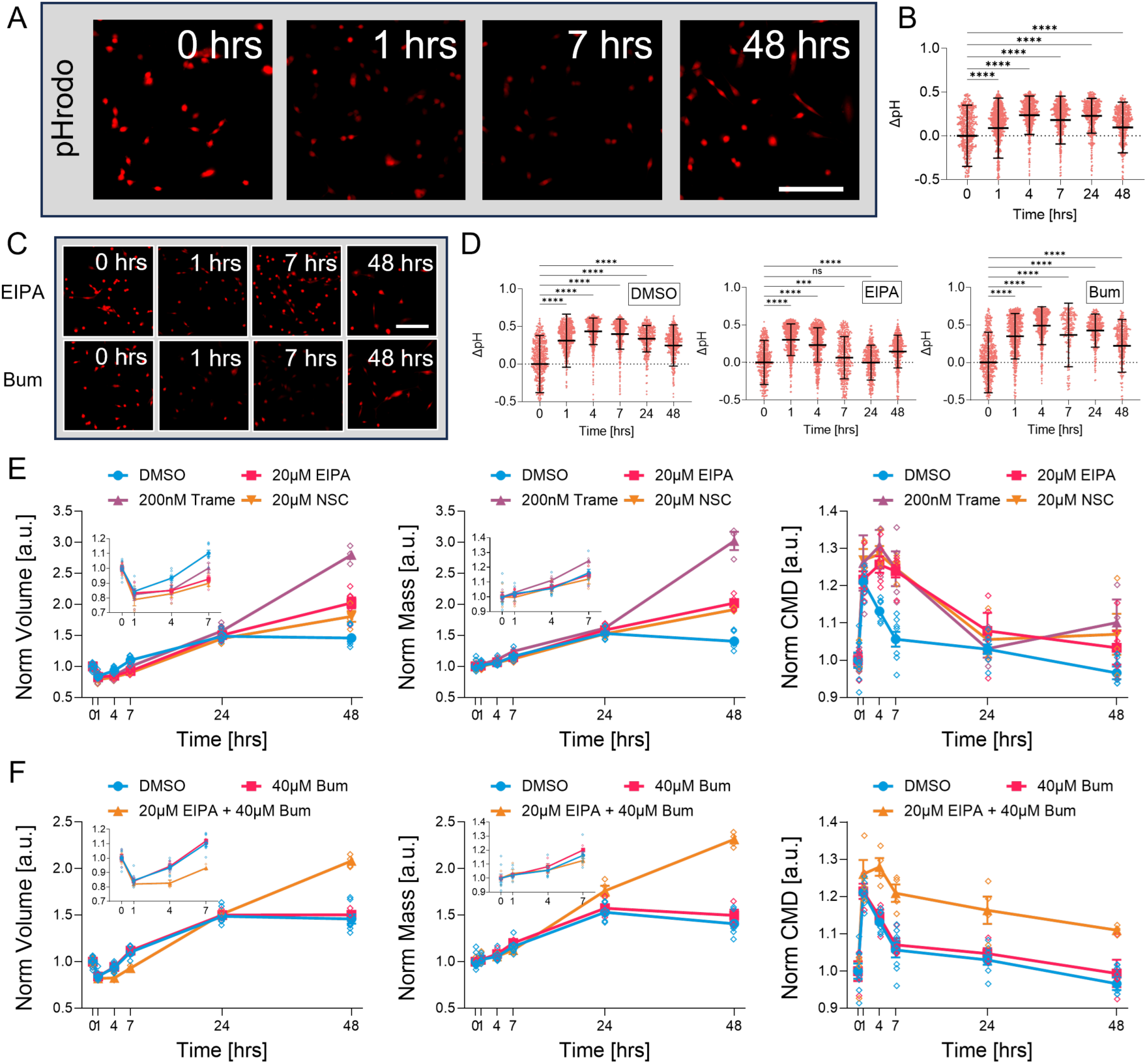
Regulatory volume increase controls density recovery. (A) Representative images of 3T3 cells with pHrodo in isotonic and 1, 7, 48 hrs during hypertonic conditions. (B) Cytoplasmic pH over time during hypertonic conditions. n ≥ 455. (C) Representative images of 3T3-pHrodo treated with 20μM EIPA and 40μM Bumetanide (Bum). (D) Cytoplasmic pH over time during hypertonic conditions treated with vehicle (DMSO), 20μM EIPA, and 40μM Bumetanide. n ≥ 207. (E) Cell volume, mass, and density over time during hypertonic conditions treated with DMSO, 20μM EIPA, 200nM Trametinib (Trame), and 20μM NSC668394 (NSC). N ≥ 4. Insets are magnified 0, 1, 4, 7 hrs data. (F) Cell volume, mass, and density over time during hypertonic conditions with DMSO, 40μM Bumetanide, and a combination of 20μM EIPA and 40μM Bumetanide. n ≥ 4. Insets are magnified 0, 1, 4, 7 hrs data. (B, D, E, F) Data are presented as the mean ± standard deviation. (B, D) One-way ANOVA tests followed by Dunn-Šidák’s multiple-comparison test were conducted between data sets. *p^ns^* > 0.05, *p*\* ≤ 0.05, *p*\*\* ≤ 0.01, *p*\*\*\* ≤ 0.01, *p*\*\*\*\* ≤ 0.0001. (A, C) Scale bars, 200 μm.

We then asked how alterations in NHE activity can influence CMD recovery. We used 20 µM EIPA to inhibit NHE, 100 nM Trametinib (Trame) to inhibit MEK, and 10 µM NSC668394 (NSC) to inhibit ezrin. We measured changes in the cytoplasmic pH by these inhibitors (Fig. 3C,D; Fig. s1H). No inhibition stopped the increase in cytoplasmic pH at 1 hour following the hypertonic shock. This indicates that the short-term NHE response to hypertonic shock is acute and robust. However, the cytoplasmic pH peaked at 1 hour and then decreased between 1 and 7 hours with EIPA and NSC treatment, in contrast to the peaking at 4 and 7 hours in vehicle control. With Trame, pH increased from 1 to 4 hours and decreased at 7 hours. These observations suggest that EIPA, Trame, and NSC inhibit NHE hyperactivation in the first 7 hours following hypertonic shock.

Following pH measurements, we also measured cell volume, mass, and CMD recovery while inhibiting MEK/ERK, ezrin, or NHE (Fig. 3E). Two key differences were observed. First, in hypertonic conditions, the regulatory increase in cell volume between 1 to 7 hours was significantly impeded when MEK, ezrin, or NHE were inhibited. Due to continuous cell mass growth, CMD further increased after hypertonic shock and peaked at 4 hours, which opposed density recovery. This indicates that MEK/ERK and ezrin-mediated NHE activity is crucial for CMD recovery in the first 7 hours under hypertonic shock. The similarity between ERK/MEK and Ezrin/NHE inhibition is potentially because MEK is a regulator of ezrin/NHE.^59,60^ NHE plays a similar role in CMD recovery in hypertonic conditions in HT1080 (Fig. s1I,J). Second, Trame, NSC, and EIPA treatments increased the steady-state cell mass density at 48 hours, suggesting a different CMD regulatory mechanism at longer timescales. We conclude that ERK/MEK and ezrin-mediated NHE activity is critically important for both regulatory CMD recovery at short times and steady-state density homeostasis at longer timescale.

Finally, we examined the impact of NKCC on CMD regulation. Although others have observed that Wnk condensation-mediated NKCC activation is responsible for cell regulatory volume increase in the first 15 minutes in hypertonic conditions,^31^ we did not observe any significant influence of NKCC inhibition with 40 µM Bumetanide on cell volume and CMD during density recovery (Fig. 3F). This is potentially because we conducted a stronger hypertonic shock (145% relative osmolarity). Inhibiting NKCC also did not influence the cell pH response (Fig. 3C,D). However, different from EIPA inhibition of NHE alone, treating cells with 20 µM EIPA together with 40 µM Bumetanide increased the steady-state cell mass density significantly in hyperosmotic conditions, indicating that the effects of NHE and NKCC channels are related, as suggested by modeling.^42,61^

### Protein synthesis rate decreases after hypertonic shock and assists in CMD recovery

In addition to cell volume regulation by NHE/NKCC, cells could also regulate CMD by changing cell mass accumulation. We thus used the SUnSET method to measure the protein synthesis rate (PSR) by culturing cells with puromycin for a 10-minute window and staining the amount of incorporated puromycin during that period.^62^ Puromycin is an analog of aminoacyl-tRNA and when inside cells, ribosomes incorporate puromycin into polypeptides that are being synthesized, thereby measuring PSR.

To start, we measured PSRs for cells in isotonic and at 1, 4, 7, 24, and 48 hours in hypertonic conditions (Fig. 4A,B). We observed a 40% decrease in PSR immediately after hypertonic shock. PSR gradually recovered within 7 hours and then overshot at 24 and 48 hours. The observed CMD decrease during the first 7 hours was partly due to the reduction in PSR and mass growth. The PSR overshoot at 24 and 48 hours indicated that the increased steady-state cell size, after recovering from the late G2 cell cycle arrest, in the hypertonic environment was achieved through an elevated mass growth. On the other hand, we stained for the cellular ubiquitin content as an indicator of the protein degradation rate (Fig. 4E, Fig. s2A).^20,63^ Although ubiquitin had a small decrease between 4 to 7 hours, it remained relatively constant throughout the density recovery process.

**Figure 4:**
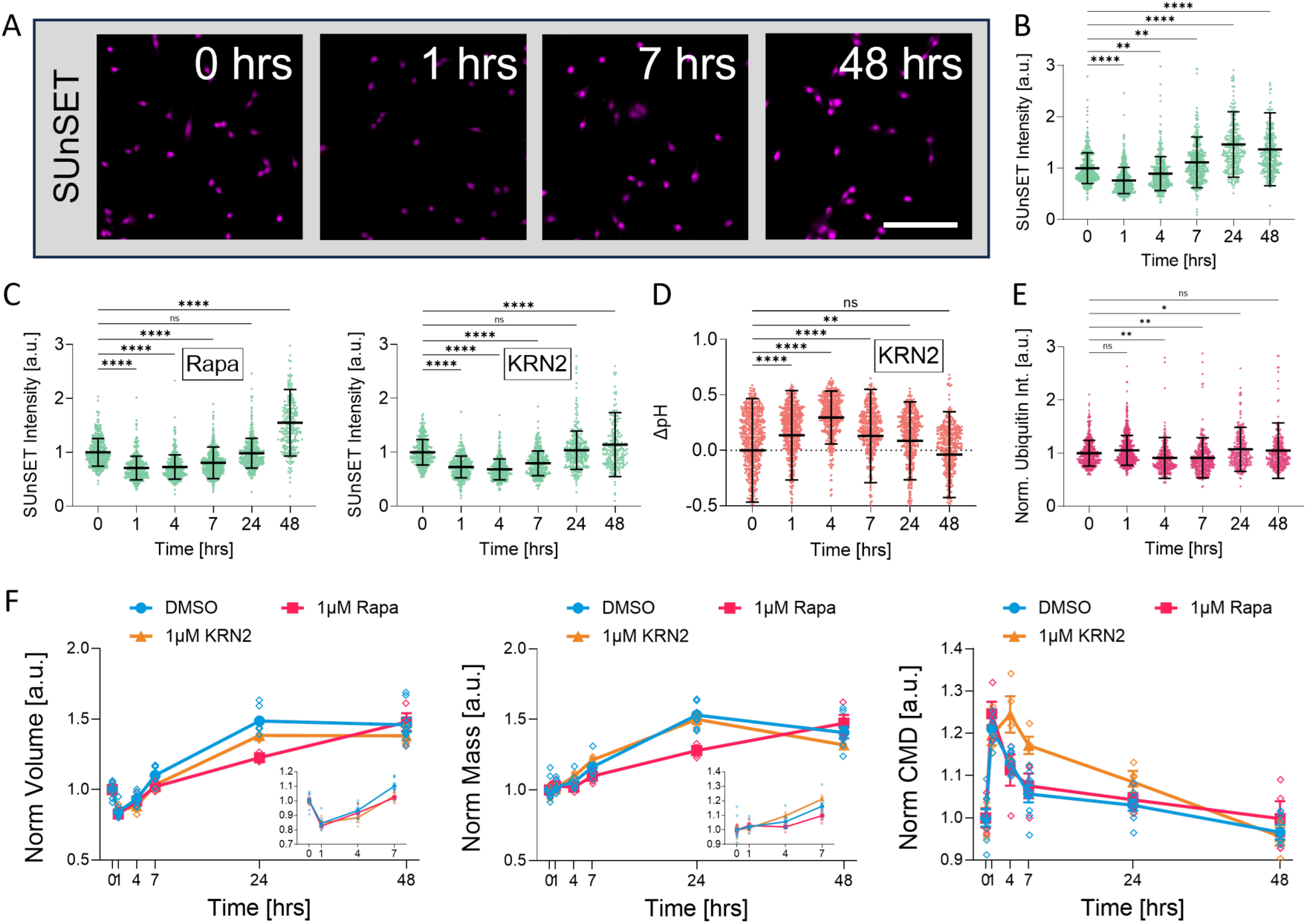
The role of protein synthesis on density recovery. (A) Representative images of anti-puromycin staining (SUnSET protein synthesis rate measurement) in isotonic and 1, 7, 48 hrs during hypertonic conditions. (B) Protein synthesis rates (PSRs) quantified using SUnSET during hypertonic conditions. n ≥ 281. (C) PSRs quantified using SUnSET in hypertonic conditions with 1µM Rapamycin and 1µM KRN2. n ≥ 241. (D) Cytoplasmic pH during hypertonic condition with 1µM KRN2. n ≥ 451. (E) Protein degradation rate quantified by immunofluorescence imaging of ubiquitin during hypertonic condition. n ≥ 281. (F) Cell volume, mass, and density plots over time during hypertonic conditions with DMSO, 1µM Rapamycin, and 1µM KRN2. N ≥ 4. Insets are magnified 0, 1, 4, 7 hrs data. (B, C, D, E, F) Data are presented as the mean ± standard deviation. (B, C, D, E) One-way ANOVA tests followed by Dunn-Šidák’s multiple-comparison test were conducted between data sets. *p^ns^* > 0.05, *p*\* ≤ 0.05, *p*\*\* ≤ 0.01, *p*\*\*\* ≤ 0.01, *p*\*\*\*\* ≤ 0.0001. (A) Scale bars, 200 μm.

Combining the protein synthesis and degradation results, we concluded that cell mass growth rate was decreased after hyperosmotic shock, which, together with cell volume dynamics, contributed to the regulatory density decrease.

We then explored the mechanism of PSR decrease. The mTOR cell growth pathway and the NFAT5-mediated osmotic stress response pathway are two candidate mechanisms.^64,65^ The mammalian target of Rapamycin complex 1 (mTORC1) is a protein complex that senses and promotes protein synthesis. Previous studies have used western blot to quantify the ratio between p-p70 and p70 and showed that mTORC1 was deactivated after hypertonic shock.^25^ The nuclear factor of activated T-cells 5 (NFAT5, TonEPB) is a transcription factor that responds to hypertonic shock by expressing osmotic stress-related genes.^27,30^ We thus inhibited mTORC1 using 1µM and NFAT5 using 1µM KRN2 and examined changes in PSR during regulatory CMD recovery (Fig. 4C). For cells with Rapamycin or KRN2 inhibition, the initial PSR decrease after the hypertonic shock was not changed, but its recovery was delayed. We also measured the cytoplasmic pH with 1uM KRN2 (Fig. 4D). The pH dynamics with NFAT5 inhibition was similar to the control, indicating that NFAT5 activity was not related to the hyperactivation of NHE.

Lastly, we measured the cell size and density over time where PSR is perturbed (Fig. 4F). We observed that mTORC1 inhibition significantly delayed the cell size overshot process but had minimal impact on regulatory CMD recovery because both volume and mass growth were slowed down. This agrees with the previous observation that inhibition of cell growth and synthesis pathways influence cell growth and proliferation but not CMD.^37^ On the other hand, after hyperosmotic shock, KRN2 inhibition only impeded the volume growth but not mass growth, which significantly reduced the density recovery. Given that KRN2 does not influence pH dynamics and NHE activity during density recovery, we hypothesized that the organic osmolytes produced following the NFAT5 signal are likely responsible for a different part of the regulatory volume growth. Organic osmolytes’ role in hypertonic stress response is believed to be secondary. Their suggested role is replacing the excess ions in the cytoplasm to achieve an electrically neutral intracellular environment.^30^ However, our experiments suggest that the organic osmolytes directly increase the cytoplasm osmolarity, influence cell volume growth, and decrease CMD.

In summary, during hyperosmotic shock-induced density increase, there is increased volume growth from hyperactivation of NHE. Cells also decrease PSR and mass growth rate in a mTORC1-independent manner. NFAT5-mediated organic osmolyte production directly mediates the volume and density recovery. Therefore, cells regulate both cell volume and mass growth during density recovery. The remaining question is what drives the reduced protein synthesis observed immediately after CMD change.

### Density-dependent N-C transport regulates protein synthesis

To explain the density-dependent protein synthesis rate change, we first exposed cells to media of different osmolarities to achieve a wider range of CMDs. We prepared two sets of media by replacing different proportions of DMEM with water plus sorbitol while maintaining consistent FBS and nutrient concentrations (Materials and Methods). Before experiments, cells were grown in the isotonic environment with the corresponding FBS and nutrient level for 24 hours. We measured the cell volume and cell mass density 1 hour after changing osmolarities (Fig. 5A). As the environmental osmolarity increased, the cell volume decreased monotonically, as reported previously.^22,66,67^ At such a short time scale, the cell mass essentially remained the same, and CMD increased monotonically due to cell volume change. In this manner, we generated CMD from 50 fg/µm³ to 300 fg/µm³ with consistent FBS and nutrient levels.

**Figure 5:**
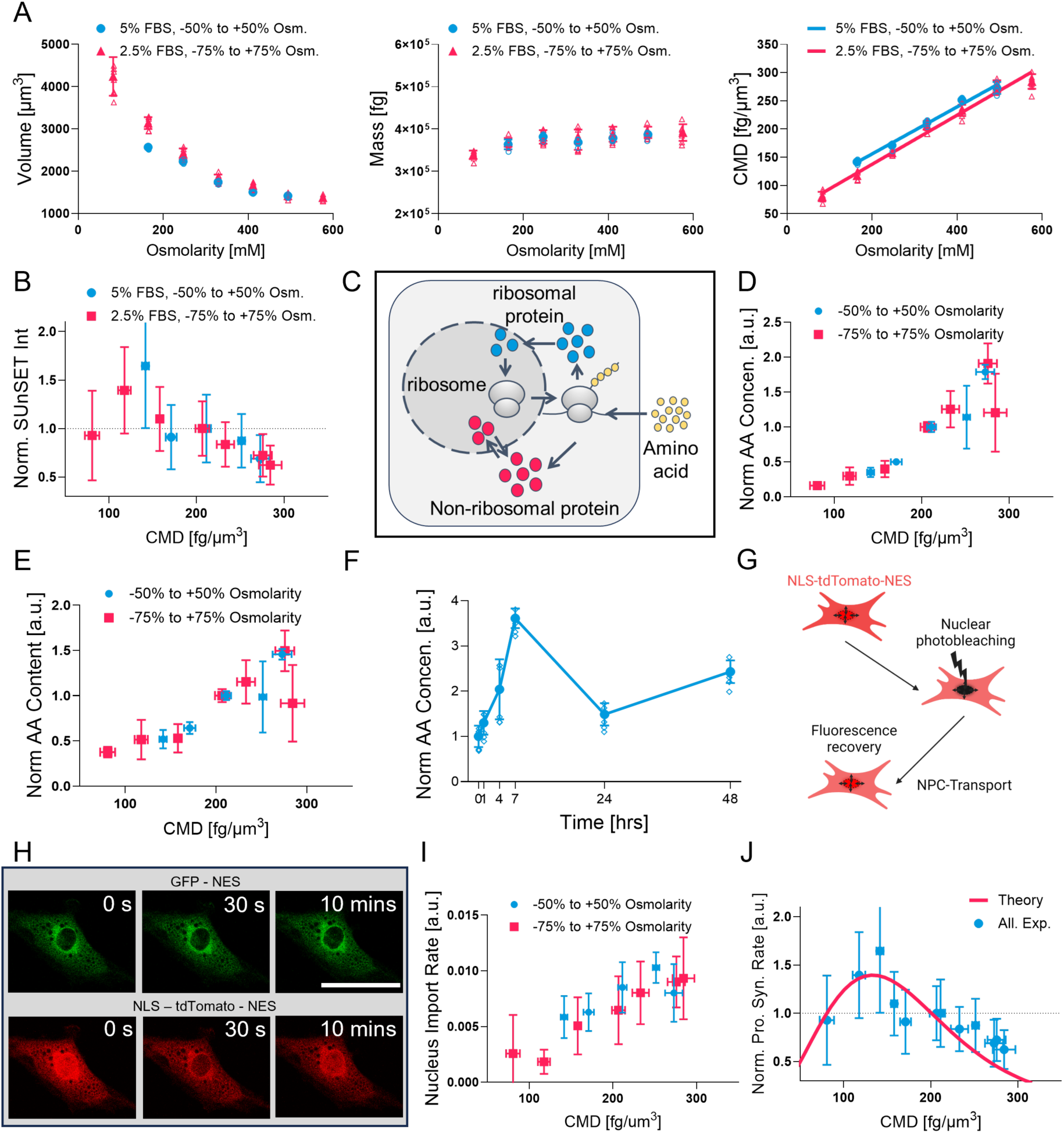
Nucleus protein import regulates protein synthesis and density recovery. (A) Cell volume, mass, and density for 1 hour in different osmotic conditions. N = 4. (B) Quantified SUnSET PSR data for different cell densities. n ≥ 212. (C) A schematic of the cell growth process involving synthesis and nucleoplasm-cytoplasm transport. (D) Cell amino acid concentration at different CMDs. N = 6. (E) Total intracellular amino acid content at different CMDs. N = 6. (F) Cell amino acid concentration over time in hypertonic condition. N = 6. (G) A schematic of the cell nucleus FRAP experiment. NLS- tdTomato-NES was photobleached to measure the protein transport rate, and GFP-NES was used as a reference to mask the nucleus. (H) Representative images of GFP-NES and NLS-tdTomato-NES before the photobleaching, at 30 s right after the nucleus NLS-tdTomato-NES photobleaching, and at 10 mins when the cell reaches a new steady state. (I) Net nucleus NLS-tdTomato-NES import rate after the nucleus NLS- tdTomato-NES photobleaching. n ≥ 9. (J) Theoretical fitting to experimentally measured protein synthesis rate as a function of density. (A, B, D, E, F, I, J) Data are presented as the mean ± standard deviation. (H) Scale bars, 50 μm.

Intuitively, as CMD increases, intracellular biochemical reaction rates should also increase due to higher concentration of all molecules. Since protein synthesis involves the binding of amino acids, mRNA, etc, to ribosomes, we expect the protein synthesis to increase with increasing cytoplasmic density. Instead, Fig. 4B showed that PSR decreased after hypertonic shock. To obtain a global view, we measured PSRs at varying CMDs by performing SUnSET experiments 1 hour after exposure to different osmolarities (Fig. 5B). We found that PSR increased as cell mass density increased for CMD lower than 147 fg/µm³. When CMD exceeded this threshold, PSR decreased as the density increased. In normal isotonic conditions, the density is around 200 fg/µm³. In this regime PSR was inversely correlated with cell mass density despite the increase of reactant concentration. This anti-correlation is a negative feedback mechanism and contributes to density regulation.

To understand the observed negative correlation between PSR and cell mass density, we can utilize simple kinetic considerations of intracellular transport, protein synthesis, and cell growth (Fig. 5C).^18^ In cells, mature cytoplasmic ribosomes bind amino acids-tRNA complex to produce polypeptides, which later mature into proteins. According to simplified chemical kinetics, 𝑃𝑆𝑅 = 𝑠 * 𝑅 * [𝐴𝐴], where 𝑠, 𝑅, [𝐴𝐴] are the synthesis rate constant, the cytoplasmic ribosome number, and the intracellular amino acid concentration, respectively. Mature ribosomes are produced by ribosome biogenesis involving nuclear the import and export of ribosomal proteins and mature ribosomes.^68,69^ During CMD change, all three quantities, 𝑠, 𝑅, [𝐴𝐴], could be changing. To reveal these details, we performed additional measurements.

To quantify the free amino acid (AA) content in cells, we used the colorimetric ninhydrin method (Materials and Methods).^70,71^ We observed that both the intracellular AA content and AA concentration increased as CMD increased 1 hour after media change (Fig. 5D,E). Combined with the decreased PSR as CMD increases, we conclude that AA consumption has decreased, which caused cytoplasmic AA content to increase. We further confirmed this by measuring cytoplasmic AA concentration 1 to 7 hours after hypertonic shock. AA also increased initially but recovered after 24 and 48 hours during CMD recovery (Fig. 5F, Fig. s2B). Together, these results suggested that AA was not the cause of PSR decrease.

Rather, the cytoplasmic ribosome number and ribosome activity, which can also change PSR, were likely the explanation for PSR decrease.

The cytoplasmic ribosome number is difficult to obtain, but since mature ribosomes are transported out of the nucleoplasm after assembly in the nucleolus, changes in the cytoplasmic ribosome number could come from changes in nucleoplasm-cytoplasm (N-C) transport. To quantify how CMD influences N-C transport, we stably expressed NLS-tdTomato-NES in 3T3 cells and used fluorescence recovery after photobleaching (FRAP) to quantify N-C transport (Fig. 5G,H).^72^ We used NES-GFP to mask the nucleus and photobleached nuclear tdTomato. The intensity recovery trajectories are shown in Fig. s2C. The net nuclear import rate and the nucleus-to-cell tdTomato ratio are shown in Fig. 5I and Fig s2D. Both results indicated that the N-C transport activity increased with increasing cytoplasmic density. Since mature ribosome complexes must be transported from the nucleus into the cytoplasm after assembly and before they can translate proteins, we conclude that CMD changes N-C transport and affects cytoplasmic ribosome number.

These experimental results can be combined with a previously developed kinetic model of protein synthesis and cell growth.^18^ This model explicitly considers the transport of ribosomal proteins, non-ribosomal proteins, and the mature ribosome complex to and from the nucleoplasm. The model also incorporates the synthesis activity of ribosomes to make new proteins and includes the scaling effects of nuclear and cytoplasmic volumes. Details of the model are given in the Materials and Methods. When we explicitly included the observed density-dependent nuclear-cytoplasmic transport, the model indeed showed that the cytoplasmic ribosome number decreased with increasing CMD (Fig. s2E), leading to a lower protein synthesis (Fig. 5J). This prediction agrees with previous data, which showed that hypertonic stress caused nuclear depletion and cytoplasm accumulation of Ran in 1 hr.^73^ This disruption on the Ran gradient would perturb the N-C transport of mature ribosome complexes.^73–79^ Changes in protein synthesis with CMD can also be explained by the synthesis coefficient, 𝑠, which may be affected by cell signaling pathways. However, our modeling suggests that this is not necessary to explain the observed PSR regulation as a function of CMD. The model also makes other interesting predictions. For instance, we found that PSR is a strong function of CMD and nuclear mass density (NMD) (Fig. s2F). In normal conditions, CMD is very close to NMD, and in this region, there is rapid change in PSR. This means that if the cell can control the cytoplasmic and/or nuclear density, it has a direct method to control protein synthesis and cell growth.

### NHE can regulate protein synthesis

Finally, we asked why 3T3 cells with NHE inhibition have a significant size overshoot 48 hours after hypertonic exposure. We first measured PSR in hypertonic conditions with NHE inhibition (Fig. 6B,C, Fig. s2G) and found that PSR increased with 20mM EIPA at 24- and 48- hour time points, suggesting that NHE inhibition increased the protein synthesis rate. We further verified that in both isotonic and hypertonic environments, 48 hours of cell culture with EIPA increased the cell protein synthesis rate, size, and density in a dosage-dependent manner (Fig. 6D,E). Therefore, NHE activity had a longer-term negative effect on protein synthesis. Together with results in Fig. 3, we found that NHE generated both cell volume increase (swelling) and regulated protein synthesis. NHE directly interacts with growth signaling molecules such as PI3K and Akt,^34,80–82^ which ultimately coordinates both cell size regulation and protein synthesis. NHE inhibition also generated HT1080 density increase in hypertonic environments, and cell size and density increase in the isotonic environment (Fig. s2H). Combining results for 3T3 and HT1080 cells, we conclude that NHE activity not only increases intracellular pH, swells cell volume, and decreases cell mass density, it also changes cell PSR depending on conditions.

**Figure 6:**
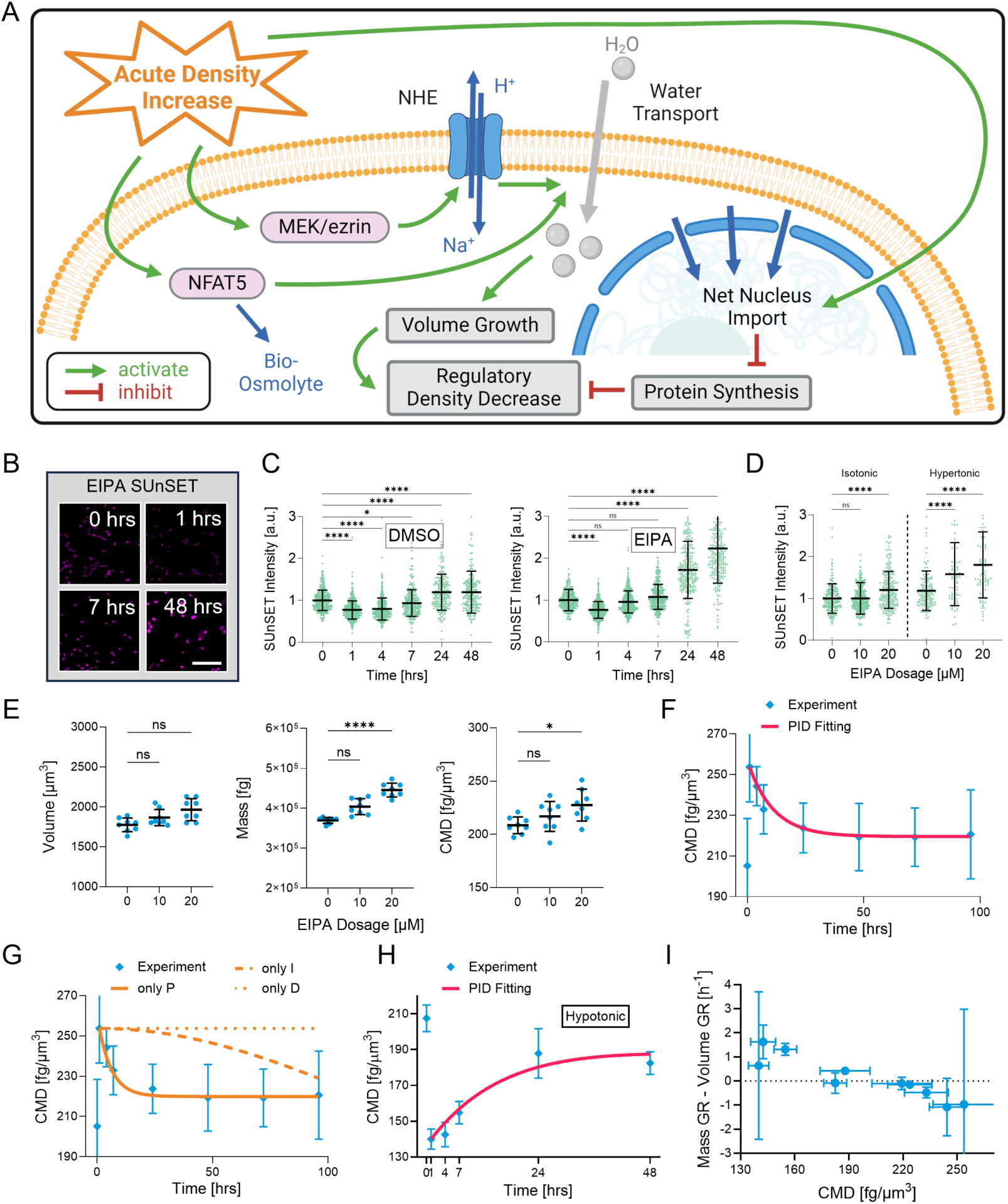
Cell mass density recovery explained by a PID control model and NHE-dependent cell growth. (A) A schematic of the CMD regulation system. (B) Original SUnSET images in isotonic and 1, 7, 48 hrs in hypertonic condition treated with 20µM EIPA. (C) Quantified SUnSET intensity over time in hypertonic conditions treated with DMSO and 20µM EIPA. n ≥ 269. (D) Quantified SUnSET intensities after 48 hours in isotonic and hypertonic conditions with different dosages of EIPA. n ≥ 29. (E) Cell volume, mass, and density in an isotonic environment treated with 20 and 40 µM of EIPA for 48 hrs. N = 8. (F) The PID control model can fit the CMD recovery curve in the hyperosmotic condition. (G) PID control model with only one proportional, integral, and differential term. (H) The PID control model also fits the CMD recovery curve in hypoosmotic (50%) conditions. (I) The difference between mass change rate and volume change rate plotted as a function of CMD in hypoosmotic and hyperosmotic conditions. N ≥ 4. (C, D, E, F, G, H, I) Data are presented as the mean ± standard deviation. (C, D, E) One-way ANOVA tests followed by Dunn-Šidák’s multiple-comparison test were conducted between data sets. *p^ns^* > 0.05, *p*\* ≤ 0.05, *p*\*\* ≤ 0.01, *p*\*\*\* ≤ 0.01, *p*\*\*\*\* ≤ 0.0001. (D) Scale bars, 200 μm.

### Cell mass density regulation can be explained by a PID model

While the observed cell volume and growth dynamics after osmotic shock are relatively complex and involve many cell components, the mass density change as a function of time can be explained by a simple control mechanism. The proportional-integral-derivative (PID) controller is a linear feedback control algorithm that can maintain system homeostasis.

While the molecular mechanism of density homeostasis is unclear, we found that the PID control algorithm could explain the density recovery curve (Fig. 6F). The target cell mass density was set to the final density at 48 hours. We further assessed the relative importance of the proportional, integral, and derivative terms in the PID controller. The proportional term had the most significant impact, while the integral and derivative terms had minimal effects (Fig. 6G, Fig. s2I). This result suggests that cells sense and adjust their density primarily by the current error in CMD, but do not integrate the prior errors or anticipate future error trends. The feedback control system relies on the current error in CMD, or concentration of proteins and possibly ions, to adjust volume and protein synthesis to reach a final target density.

We note that the CMD regulatory recovery is also observed for hypotonic shock (Fig. 6H, Fig. s2J). Again, PID control is sufficient to explain CMD recovery. In both hypotonic and hypertonic conditions, cells adjust the rate of cell volume change and the rate of cell mass change to achieve CMD homeostasis. Since at steady state, the rate of cell volume change and the rate of cell mass change as a function of CMD must be equal, plotting the difference between these rates vs CMD gives the final homeostatic CMD (Fig. 6I). The homeostatic CMD is a weak function of external media osmolarity.

## Discussion

Recent studies demonstrated that the cell mass density is robustly maintained in different cell types. This study demonstrates how CMD is regulated in mammalian cells after a hypertonic shock. We first show that CMD is conserved in several different tissue cell types. We then show that CMD increases immediately after exposure to hyperosmotic shock and gradually recovers over 48 hours, and this regulatory density recovery process is cell cycle independent. We discover that NHE activation generated volume increase and protein synthesis decreased during CMD recovery. The decreased protein synthesis rate is effectively a negative feedback element in controlling CMD. Moreover, we show that the nuclear-cytoplasmic transport can regulate the protein synthesis rate and CMD recovery. Taken together, the results suggest that cells achieve density regulation by sensing the difference between the current density and a target density.

Several remaining questions deserve future exploration: 1) Is there a target CMD that cells like to maintain? How do cells sense the error between current and target densities? We hypothesize that the density error can be sensed from the concentration of a specific molecule or groups of different molecules. However, the identity of these molecules or ions needs to be revealed. 2) How does NHE activity influence the protein synthesis rate? We observed that NHE inhibition increases the protein synthesis rate. Currently, the understood role of NHE is the regulation of cell pH and volume. We hypothesize that the NHE inhibition disrupts the CMD regulatory system. One of the outputs of the CMD regulatory system is protein synthesis, and any disruption of this system can potentially change PSR.

The CMD control system probably involves a large number of ion channels/transporters and their regulators, such as PI3K/Akt. Direct and indirect physical interactions between NHE and PI3K/Akt have been identified.^80^ Therefore, the CMD regulatory system potentially involves ion channels/transporters and canonical signaling pathways such as PI3K and beyond. Our results suggest that the CMD system can simultaneously regulate cell volume (by regulating ionic fluxes and cell water content) and cell mass (by regulating protein synthesis). PSR is also influenced by N-C transport and ribosome biogenesis, which naturally provides negative feedback on PSR when CMD changes. These PSR changes as a function of CMD are captured by our model of cell growth, which incorporates these elements in a simple kinetic form.

In addition to PSR regulation described by our model, the model also predicts that the CMD and the nuclear mass density (NMD) can influence PSR and the cell growth rate (Fig. s2F). In fact, the cell growth rate is a sharp function of CMD and NMD. When CMD and NMD become unequal and deviate too far from the physiological values, cell growth arrest can occur. The natural CMD and NMD sit very close to the transition region. From the point-of-view of optimal control, this suggests that the cell can easily modulate the growth rate by changing CMD and NMD. This implies that the CMD regulatory system is a master regulator of cell growth, and by changing cell water/ionic content, the cell can modulate growth. Of course, the actual details of this regulatory system are much more complex than our simple model suggests. CMD homeostasis is likely an emergent outcome of a complex cell network of interactions that integrates environmental information (osmolarity, pressure, viscosity, matrix properties, and cytokines) with the internal cell state and phenotype.

Deleterious alterations in the CMD regulatory system can potentially lead to cellular senescence and cancer-like cell states.^39,83^ Other possible states of the CMD network may be behind cell phenotype specification, and growth and morphogenesis in general.

## Acknowledgments

This work has been supported in part by NIH R01GM134542 (to S.X.S) and the Air Force OTice of Scientific Research FA9550-22-1-0334 (to I.B). We would like to thank Morgan Benson, Ana Carina Nogueira Vasconcelos, Mohammad Ikbal Choudhury, Blake Johnson, Hongyue Cui, and Rong Li for their help with experiments and discussions. We also thank the Johns Hopkins University Integrated Imaging Center for flow cytometry support, and the Johns Hopkins University WSE Whitaker Microfabrication Lab for microfabrication device support.

## Material and Methods

### Cell culture

NIH3T3 cell line was purchased from the American Type Culture Collection (ATCC). HT1080, T47D, and Hs578 cell lines were a gift from K. Konstantopoulos (Johns Hopkins University, Baltimore, MD). RPE 2n and RPE 4n cell lines were a gift from R. Li (National University of Singapore, Singapore). MCF-10A, MDA-MB-231, and its single cell clones (SCCs) MDA-SCC-1006, MDA-SCC-1304, and MDA-SCC-1308 cell lines were provided by D. Wirtz (Johns Hopkins University, Baltimore, MD). SCCs of MDA-MB-231 were generated through the expansion of individual parental cells as described in Ref.^1^ HT1080 SCCs were generated using the same method. NIH3T3, HT1080, RPE 2n, RPE 4n, T47D, Hs578, MDA-MB-231, and all SCCs were cultured in Dulbecco’s Modified Eagle Medium (DMEM; Corning) supplemented with 10% fetal bovine serum (FBS; Sigma) and 1% antibiotic solution containing 10,000 units/mL penicillin and 10,000 μg/mL streptomycin (Gibco).

MCF-10A cells were cultured in DMEM/F-12 supplemented with 5% horse serum, 20 ng/mL Epidermal Growth Factor (EGF), 0.5 μg/mL hydrocortisone, 100 ng/mL cholera toxin, 10 μg/mL insulin, and 1% antibiotic solution. Cells were passaged using 0.05% trypsin-EDTA (Gibco). All cell cultures and live cell experiments were conducted at 37°C in a humidified atmosphere with 5% CO2.

### Preparation of media with di4erent osmolarity

The hypertonic media was supplemented with D-sorbitol in DMEM (MilliporeSigma) (329 mOsm) to achieve different osmolarity. Unless specified, all hypertonic media used in this work contains 148 mM (27.9 g/L) D-Sorbitol with a final osmolarity at 477 mOsm. For hypotonic shock, 50% hypotonic media (165 mOsm) was prepared by mixing 50% ultra- pure water with 50% cell culture media. To maintain consistent nutrient concentrations in the hypotonic shock experiment, cells were pre-incubated in an isotonic solution consists of 50% cell culture media and 50% 329 mOsm sorbitol water solution for 24 hrs. The osmolarity of 10% FBS medium were measured using an Advanced Instruments Model 3320 osmometer, (Table. s1). The osmolarity of 2.5% FBS and 5% FBS medium were calculated based on the measurements on 10% FBS medium.

In selected experiments shown in Fig. 5, we also prepared a series of media with varying relative osmolarities ranging from -75% to +75 % with consistent nutrient conditions. For the first series (Media 1), all media contains 25% DMEM, 2.5% FBS, and distilled water. For the second series (Media 2), all media contains 50% DMEM, 5% FBS, and distilled water. D-sorbitol was added to adjust the final osmolarity to a range of -75% to +75 % relative to that of isotonic media for Media 1, and -50% to +50% relative osmolarity for Media 2. In all experiments, cells were pre-adapted in the corresponding isotonic media (+0%) for 24 hrs. After that, cells were exposed to media with different osmolarity for 1 hr.

### Pharmacological inhibitors

In select experiments, cells were treated with the following pharmacological agents: Vehicle control: 0.25% DMSO (Invitrogen), 20 μM EIPA (Tocris), 0.2 μM Trametinib (MedChemExpress), 0.25 µM Doxorubicin (SelleckChem), 1 µM Rapamycin (Biogems), 1 µM KRN2 (MedChemExpress), 10 µM NSC668394 (MilliporeSigma), 40 µM Bumetanide (Tocris). For all drug-related experiments, cells were pre-adapted to the pharmacological agents for 3 hrs prior to the experimental conditions.

### Fluorescence eXclusive method (FXm) device preparation

A detailed protocol can be found in our previous paper.^2^ In brief, FXm channel masks were designed using AutoCAD and ordered from FineLineImaging. Silicon molds were fabricated using SU8-3010 (Kayaku) photoresist following standard photolithography procedures and manufacturer’s protocol. One layers of photoresist were spin coated on a silicon wafer (IWS) at 500 rpm for 7 s with acceleration of 100 rpm/s followed by 2,000 rpm for 30 s with acceleration of 300 rpm/s. After a soft bake of 5 mins at 95 °C, UV light was used to etch the desired patterns from negative photoresist to yield feature heights that were 10 μm. The length of the channels is 16 mm and the width is 1.2 mm.

A 10:1 ratio of PDMS Sylgard 184 silicone elastomer and curing agent were vigorously stirred, vacuum degassed, poured onto each silicon wafer, and cured in an oven at 80 °C for 45 min. Razor blades were then used to cut the devices into the proper dimensions, and inlet and outlet ports were punched using a blunt-tipped 21-gauge needle (McMaster Carr; 76165A679). The devices were cleaned by sonicating in 100% isopropyl alcohol for 10 min, and dried using a compressed air gun. The devices and sterilized 50-mm glass-bottom Petri dishes (FlouroDish Cell Culture Dish; World Precision Instruments) were exposed to oxygen plasma for 1 min for bonding. The bonded devices were then placed in an oven at 80 °C for 45 min to further ensure bonding.

### Fluorescence eXclusion method (FXM) data analysis and cell volume calculation

Individual cells were tracked by customized MATLAB code using the algorithm described in our previous paper.^3^ In brief, individual cells were masked, and the mask were expanded by 10 pixels (∼3 μm) in each direction to ensure proper cell cropping. Any overlapping cells were discarded. After all images at different time points were processed, we ran an automatic cell tracking algorithm based on the position of the geometric center of each cell mask. The results from the automatic cell tracking algorithm were then examined manually by one of the authors. The mean fluorescence intensity of the pixels outside FXm channel 𝐼*_Noise_*, reflects the environmental fluorescence noise. The mean fluorescence intensity of the pixels outside cell masks, defined as the mean background intensity 𝐼*_Channel_*, reflects the height of the FXm channel. The local intensity map within the cell mask, 𝐼*_cell_*, reflects the difference between channel height and cell height. Given a known channel height CH and the intensity map 𝐼*_cell_*, cell volume is calculated as

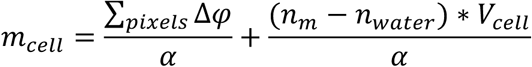

After each experiment, the channel height of all FXm devices was measured using a confocal microscope using Z-stacking. Confocal microscopy has strong point spread function that affects the precision of channel height measurement. To overcome this, we determined the point spread function using small fluorescence beads (0.2 μm, Invitrogen) and deconvoluted the intensity profile, and the channel height was obtained after the deconvolution.

### Epi-fluorescence and confocal microscopy

For epi-fluorescence imaging, a Zeiss Axio Observer inverted, wide-field microscope using a 20× air, 0.8-NA objective equipped with a Hamamatsu Flash4.0 V3 sCMOS camera was used. For confocal imaging, a Zeiss LSM 800 confocal microscope equipped with a 20× air, 0.8--NA objective was used. All microscopes were equipped with a CO2 Module S (Zeiss) and TempModule S (Zeiss) stage-top incubator (Pecon) that was set to 37 °C with 5% CO2 for live cell imaging. For imaging of immunofluorescence assays, the samples were imaged under room temperature and without CO2. ZEN 2.6 or 3.6 Software (Zeiss) was used as the acquisition software.

### Quantitative phase microscopy

Cell dry mass represents the total mass of all intracellular non-water molecules. To measure cell dry mass, we utilized the Holomonitor M4 quantitative phase microscope (PHI). Quantitative phase microscopy (QPM) is based on the principle that the refractive index of a material inversely correlates with the velocity of light passing through it. When in operation, the Holomonitor directs a laser beam through both the cell and the surrounding culture medium. As light passes through the cell, which has a higher refractive index than the medium, it slows down, resulting in a detectable phase shift. This phase shift is captured by the camera and processed to summarize the total phase alteration caused by the entire cell (optical volume). The total phase alteration of individual cells was analyzed using the dedicated Holomonitor M4 software. For aqueous solutions, the refractive index is linearly proportional to the solute mass concentration, with a consistent factor 𝛼 = 0.18 mL/g irrespective of the solute type.^4–6^ We have also validated this factor using a NaCl and a BSA solution (Fig. s1D). After obtaining the cell volume, the optical volume, and culture medium refractive index, cell mass can be calculated as:

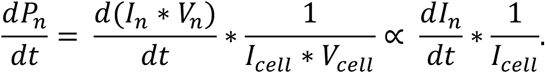

where Δ𝜑 is the phase shift cells make in each pixel, n*_m_* is refractive index of cell culture medium, 𝑛*_water_* is refractive index of water. The refractive index of different media was measured using the Digital Brix Refractometer (MILWAUKEE Instruments MA871; Table s1, s2, s3).

### Cell mass density (CMD) measurements

For population-level cell volume, mass and CMD measurements, the average volume and optical volume for a population of cells were first measured using FXm and QPM. Mass and CMD were calculated based on the population average of cell volume and optical volume as described above. For each experiment, an equal number of cells from the same biological parent were seeded into T25 flasks, with one flask assigned to each condition.

For the osmotic shock experiments, the medium in each flask was replaced with hypotonic or hypertonic media 4-96 hrs before imaging. Cells were then seeded into collagen coated FXm channels for volume measurement or lumox 24-well plate (Sarstedt) for optical volume measurement 1hr before imaging. For 1 hr exposure conditions, cells were seeded directly with the osmotic shock media. For all drug-related experiments, cells were pre-adapted to the pharmacological agents for 3 hrs prior to the osmotic shock. All conditions were imaged at the same time. Cell volume and optical volume at each time point were always measured in parallel from the same flask of cells to ensure consistency.

For FXm experiment, cells were centrifuged and concentrated to 2 × 10⁶ cells/mL, and seeded into FXm channel with 0.2 mg/mL Alexa Fluor 488 Dextran (MW 2,000 kDa; ThermoFisher) dissolved in corresponding media. For QPM, 1.5 × 10⁴ cells were seeded into each well of 24-well plates. In all experiments, microfluidic devices or plates were incubated with 50 μg/mL of type I rat-tail collagen (Enzo) for 1 hour at 37°C. Following incubation, the devices were thoroughly washed with the corresponding media. In selected experiments Fig. 2C insets, cell volume and optical volume were tracked over time at a single cell level. In these experiments, cells were prepared as described above in isotonic media and were seeded into FXm channels or 24 well plates for three hrs. Following attachment, cells were imaged at five-minute intervals for 30 mins in isotonic media. Then, osmotic shock were applied by injecting the corresponding media into the FXm channels or switching the media in the 24-well plates. Imaging continued for an additional hour, capturing the same cells throughout the process. Volume and optical volume were tracked over time for individual cells as described above. The population average of cell volume and optical volume were then used to calculate cell mass and density.

### Tomocube Holotomography microscopy and single cell volume, mass, and mass density (CMD) measurements

For single cell mass, volume, and CMD measurements, the 3D quantitative phase images were acquired using an interferometry-based tomographic microscope, HT-2H (TomoCube). To maintain optimal conditions during image acquisition, an in-situ incubator, TomoChamber (TomoCube), was used to ensure a stable temperature of 37°C, 5% CO2 concentration, and a humid atmosphere. The microscope employed a 532 nm continuous-wave laser, a 60x water immersion objective (NA = 1.2), and a digital micromirror device.

The resulting 3D refractive index (RI) tomograms provided a lateral resolution of 95 nm and an axial resolution of 190 nm. For 3D segmentation and analysis, cells were initially segmented in 2D using the maximum intensity projection of the tomogram, utilizing custom-developed code based on our previous paper.^7^ These 2D segments were then applied as masks across all slices in the tomogram to isolate individual cells in each field of view. Subsequent 3D segmentation was performed using a threshold refractive index of 1.36 for NIH3T3 cells and 1.355 for HT1080 cells. Cell volume was determined by counting the number of non-zero pixels in the segmented image, while cell optical volume was calculated by integrating the pixel size multiplied by the pixel refractive index above the background for all non-zero pixels.

In all tomographic experiments, cells were seeded into collagen-coated 1.5H glass coverslip bottoms of TomoDishes (TomoCube) at a density of 3 × 10⁴ cells and allowed to attach overnight. Following attachment, cells in different dishes were exposed to hypertonic medium for the corresponding durations as of the population measurement. Imaging of all samples was conducted on the third day after seeding, using TomoCube Holotomography technology.

### Intracellular pH measurements using pHrodo

In all pH experiments, cells were seeded at a density of 3,000 cells per well into glass bottom 24 well plate (Cellvis) with collagen coated as described above two days prior to the experiments. The osmotic shock media were switched corresponding to the condition 1 hrs to 48 hrs before measurement, and all conditions were imaged at the same time.

Intracellular pH was measured using pHrodo Red AM (Invitrogen, P35372), following the manufacturer’s instructions. In brief, cells were incubated with 1 mL of media containing a dilution of 1 μL of 5 mM pHrodo Red AM in 10 μL of PowerLoad 100X concentrate at 37°C for 45 mins. After incubation, the cells were gently washed once with fresh media and allowed to settle for 15 mins inside the microscope incubator before imaging. To obtain the absolute pH of different cell types, an intracellular pH calibration buffer Kit (Invitrogen, P35379) was used with pHrodo Red AM, following the manufacturer’s protocol. The pHrodo intensities from individual cells, measured above the background, were used to calculate the intracellular pH of each cell.

### SUnSET protein synthesis rate measurements

The SUnSET method was employed to measure the single-cell protein synthesis rate. Cells were seeded and prepared as described above in the pH section. Before fixation, cells were treated with 100 µg/mL puromycin (Sigma-Aldrich, P8833) diluted in the corresponding osmotic shock media for 10 mins in the incubator. Following puromycin treatment, cells were immediately fixed using 4% paraformaldehyde (PFA) for 10 mins, permeabilized using 0.1% Triton on ice for 10 mins, and blocked using a blocking buffer containing 3% FBS, 3% BSA, and 3% goat serum in PBS solution at RT for 1 hr. Cells were washed using PBS for 3 times between each step. Cells were then stained with anti-puromycin conjugated antibody (1:200; EMD Millipore, MABE343, clone 12D10) in 1% FBS, BSA, and goat serum PBS solution overnight at 4°C, followed by another 3x wash. All samples were imaged using an epi-fluorescence microscope to obtain the total intensity.

### Propidium iodide DNA content measurements

In all experiments, cells were seeded at a density of 6 x 10^4^ cells per T25 flask two days prior to the experiments. The hyperosmotic media were switched corresponding to the condition 1 hrs to 48 hrs. Cells from all conditions were then detached using Trypsin at the same time, concentrated by centrifugation, and fixed using ethanol at 4°C overnight. After fixation, the cell pellet was gently washed once with PBS, re-centrifuged, and stained with 1ml 1.0 mg/mL propidium iodide solution (Invitrogen, P3566) every 1 x 10^6^ cells for 45 mins. The propidium iodide intensities from individual cells measured by Flow Cytometer were considered as the cell DNA content.

### FRAP nuclear transport rate measurements and transfection

To investigate the nuclear transport rate, we first generated NIH3T3 cell line stably co-expressing NES-GFP (Addgene, 17301) and NES-tdTomato-NLS (Addgene, 112579) as described below. The fluorescence recovery after photobleaching (FRAP) experiment was performed on a Zeiss LSM800 confocal microscope (Zeiss) with 40X oil immersed objective and the bleaching module. Cells were seeded and treated as described above before experiment. To bleach cell nucleus NES-tdTomato-NLS, we first drew a mask of cell nucleus based on the NES-GFP signal, then we bleached the cell nucleus NES-tdTomato-NLS with 561nm laser at 20% intensity for 100 iterations. After bleaching, cells were imaged continuously for 10 mins with 10-second frame rate. To analyze the nucleus intensity recovery, we first subtracted the image background intensity and drew a rectangular box inside the nucleus based on the NES-GFP images. After that, we calculated the average NES-tdTomato-NLS intensity inside the box.

We calculated the nucleus tdTomato’s intensity change after normalizing to cells’ total fluorescence content right after photobleaching as the net nuclear protein import rate with the equation:

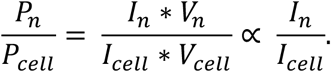

Here 𝑡, 𝑃_n_, 𝐼_n_, 𝐼_cell_, 𝑉_n_, 𝑉_cell_ are time, nucleus protein number, mean nucleus tdTomato intensity, mean cell tdTomato intensity, nucleus volume, and cell volume, respectively. We assumed 𝑉_n_ / 𝑉_cell_ is a constant during the transport experiment.

We used the last image of the FRAP video (the image at 10 mins) to analyze the steady-state nucleus-to-cell NES-tdTomato-NLS content ratio. We first subtracted the image background, then we masked the cell and nucleus based on the NES-GFP image. The nucleus-to-cell NES-tdTomato-NLS ratio was calculated as the average nucleus tdTomato intensity divided by the average whole cell tdTomato intensity. We used the nucleus-to-cell intensity ratio as an approximation of the nucleus to cell protein content based on the equation:

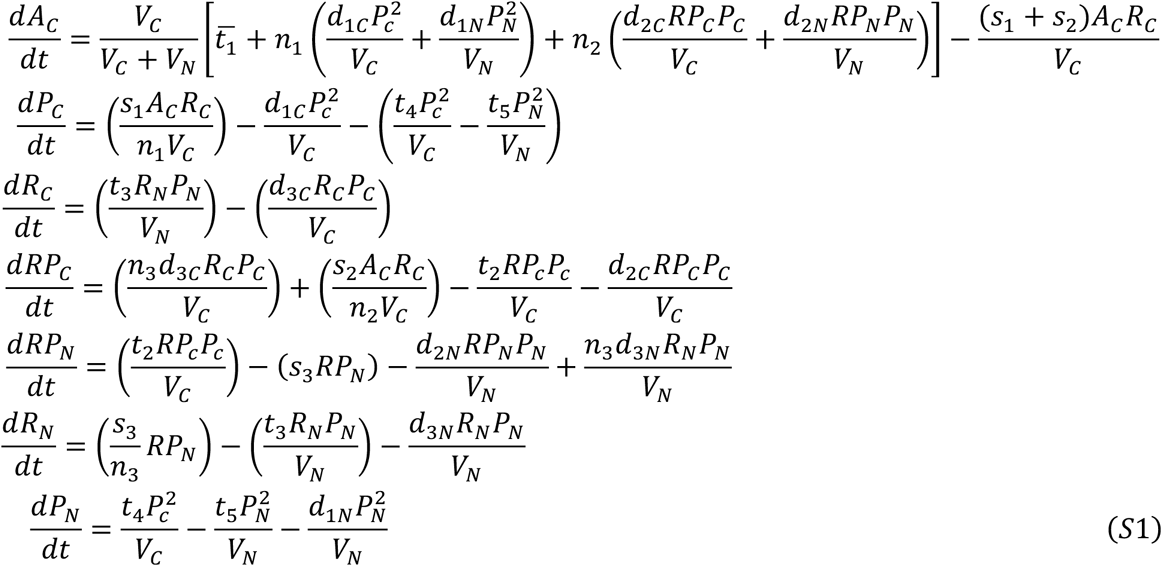

### Colorimetric ninhydrin method

The colorimetric ninhydrin method was performed to measure intracellular amino acid content. We seeded 3 × 10^4^ cells in tissue culture treated plastic bottom 6-well plates 72 hrs before experiments. We always prepared two sets of cells for each biological repeat, and the two sets of cells were treated in different osmotic shock as described above simultaneously. Before amino acid measurement, one set of the cells were counted to determine the change in cell number due to treatment. We then lysed the other set with 300 μL 80% methanol (pre-cooled to -20 °C) for 10 mins on ice. All samples were then scraped and collected into 1.5 mL tube, followed by centrifugation at 18,000 g for 10 mins at 4 °C. After that, 3 x 50 μl supernatant were added into a 96 well plate to generate 3 technical repeats for each condition. Samples were then mixed with 2% Ninhydrin (Sigma; dissolved in ethanol) at a 1:1 ratio and heated in a 95 °C oven for 10 mins to facilitate the reaction. The results were then read using a colorimeter (Molecular Devices) at 570 nm absorbance. An amino acid standard was also generated in the same plate using L-alanine solution with a concentration gradient from 0 to 250 μg/mL. The cell amino acid content was obtained by calibrating the sample readouts against the standard curve, and adjust based on the cell number in each condition. Amino acid concentrations were calculated by dividing the total amino acid content by average cell volume measured from FXm.

### FUCCI Cell Cycle Length Quantification

We expressed RFP-cdt1 and GFP-Geminin in NIH3T3 using Premo FUCCI cell cycle sensor (Invitrogen) following manufacturer’s protocol. In experiments, cells were seeded as described in the pH section. We imaged cells immediately after the hypertonic shock using Epi-fluorescence microscopy for 60 hrs with 15 mins frame rate. Cell G1 and G2 length were tracked on the first generation of daughter cells born after hypertonic shock by visually examining cell nucleus RFP and GFP intensity.

### Live cell reporters, lentivirus preparation, transduction, and transfection

For lentivirus production, HEK 293T/17 cells were co-transfected with psPAX2, VSVG, and the lentiviral plasmid of interest. 48 h after transfection, the lentivirus was harvested and concentrated using centrifugation. NIH3T3 cells with inducible NES-GFP expression was generated from our previous work.^8^ To generated stable expression of NES-tdTomato-NLS (Addgene, 112579), NIH3T3 NES-GFP cells at 60–80% confluency were incubated for 24 h with 100X virus suspension and 8 μg/mL of Polybrene Transfection Reagent (Millipore Sigma). Cells were then selected for NES-tdTomato-NLS signal using flow cytometry.

Expression of NES-GFP was induced by doxycycline (1 μg/mL) for 16 h before experiment.

### Model description

Cell mass/volume increase can be described by a minimal eukaryotic cell growth model, which includes protein synthesis, transport and degradation. The dynamical equations are^9^:

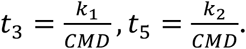

Here 𝐴_c_, 𝑃_c,N_, 𝑅_c,N_, 𝑅𝑃_C,N_ are cytoplasmic/nucleoplasmic numbers of amino acids, non-ribosomal proteins, ribosomes, and ribosomal proteins, respectively. 𝑛_1_, 𝑛_2_ are average amino acid numbers per non-ribosomal and ribosomal protein. 𝑛_3_ is the average number of proteins in a ribosome. 𝑡̅_1_ is the amino acid uptake rate, which includes the exterior amino acid abundance, transport efficiency and percentage of transport proteins. 𝑡_2_, 𝑡_3_, 𝑡_4_, 𝑡_5_ are transport coefficients of 𝑅𝑃_c_, 𝑅_N_, 𝑃_c_, 𝑃_N_, which include transport efficiency and percentage of transported proteins. 𝑑_1c_, 𝑑_2c_, 𝑑_2N_, 𝑑_3c_, 𝑑_3c_, 𝑑_3N_ are degradation or disassembly coefficients, which include degradation efficiency and percentage of degradation (assisting) proteins. 𝑠_1_ and 𝑠_2_ are synthesis coefficients of non-ribosomal and ribosomal protein, which include mRNA concentration and ribosome moving speed. 𝑠_3_ is the assembly coefficient of ribosome, which includes the number of assembly sites and assembly rate at each site. 𝑉_N_ and 𝑉_c_ are nucleus and cytoplasm volumes, respectively. Notably, the terms describing nuclear export have been modified. In Ref,^9^ it was assumed that both import and export rates across the nuclear envelope are determined by the number of nuclear pore complexes (NPCs), which is considered proportional to the cytoplasmic protein count (𝑃_c_). This is a reasonable assumption for long term behavior during the time scale of cell cycles. However, in this paper, we are describing growth change within an hour. Consequently, we assumed that the number of NPCs remains constant, and the export rate depends on the concentration of RanGTP inside the nucleus,^10^ which is proportional to the nuclear protein number (𝑃_N_). The modified terms include ribosome export rate 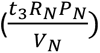 and the non-ribosomal protein export rate 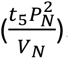 Based on experimental findings indicating an increase in net nuclear import with higher CMD, we also assumed that the nuclear export coefficients (𝑡_3_, 𝑡_5_) are decreasing functions of cell mass density. Specifically, 𝑡_3_, 𝑡_5_ were assumed to be inversely proportional to CMD: 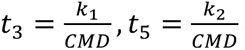.

𝑉_c_ and 𝑉_N_ are related to total macromolecule number via number densities 𝑟_1_ and 𝑟_2_ (which are proportional to mass densities):

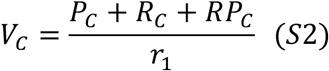

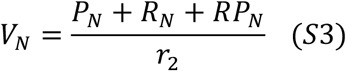

We can tune cytoplasmic and nuclear density and examine how they influence cell mass regulation via protein synthesis.

### Exploring the influence of mass density on the protein synthesis rate

We first simulated the density-dependent protein synthesis rate with our model. In this simulation, we assumed that nuclear and cytoplasmic densities are equal (𝑟_1_ = 𝑟_2_). The nucleus protein transport rate is in the order of 5 * 10^6>^𝑠^-2^, and the cell membrane amino acid transporter turnover rate is in the order of 100𝑠^-2^,^11^ which is much smaller than the time scale (1 hr) for our experiment. On the other hand, the density recovery process takes 48 hrs which is larger than the time scale for our experiment. Considering the time scales, we also assumed that cells were observed under a semi-steady protein synthesis rate state under corresponding densities.

We then varied the number density (𝑟_1_, 𝑟_2_) to see how it influences protein synthesis rate (including both ribosomal and non-ribosomal proteins), which is defined as:

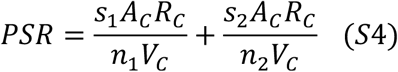

We solved equation (S1) with varying number/mass densities (𝑟_1_, 𝑟_2_), while maintaining the same cytoplasm and nucleus number/mass density (𝑟_1_ = 𝑟_2_), and calculated the corresponding steady-state protein synthesis rate and the normalized cytoplasmic ribosome number (Fig. 5J, Fig. s2E). Finally, we also simulated the cell growth rate when varying both the cytoplasm and nucleus density (Fig. s2F). All parameters were fitted to the experimental data using the random search method. The fitted parameters are as follows:𝑡_1_ = 101.73, 𝑡_2_ = 0.91, 𝑡_3_ = 0.097, 𝑡_4_ = 0.019, 𝑡_5_ = 0.2436, 𝑠_1_ = 38.18, 𝑠_2_ = 28.6987, 𝑠_3_ = 1.5033. The proportional coefficients of transport coefficients with respect to the 1/CMD are 𝑘_1_ = 2.9, 𝑘_2_ = 0.6.

### Quantification and statistical analysis

In this study, at least two independent biological repeats were conducted for every experimental condition. The number of cell groups (N) or single cells (n) analyzed for each specific experimental condition was noted in the figure captions. All data in this article represent the mean ± SD of pooled data from all experiments. Statistical analysis was conducted using GraphPad Prism 9 or 10 (GraphPad Software). One-way ANOVA tests followed by Dunn-Šidák’s multiple-comparison test were conducted between data sets. *p*^ns^ > 0.05, *p*\* ≤ 0.05, *p*\*\* ≤ 0.01, *p*\*\*\* ≤ 0.01, *p*\*\*\*\* ≤ 0.0001. The statistical analysis of all cell mass density over time during hypertonic condition experiments is shown in Fig. s3.

---

**Supplemental Figure 1:**
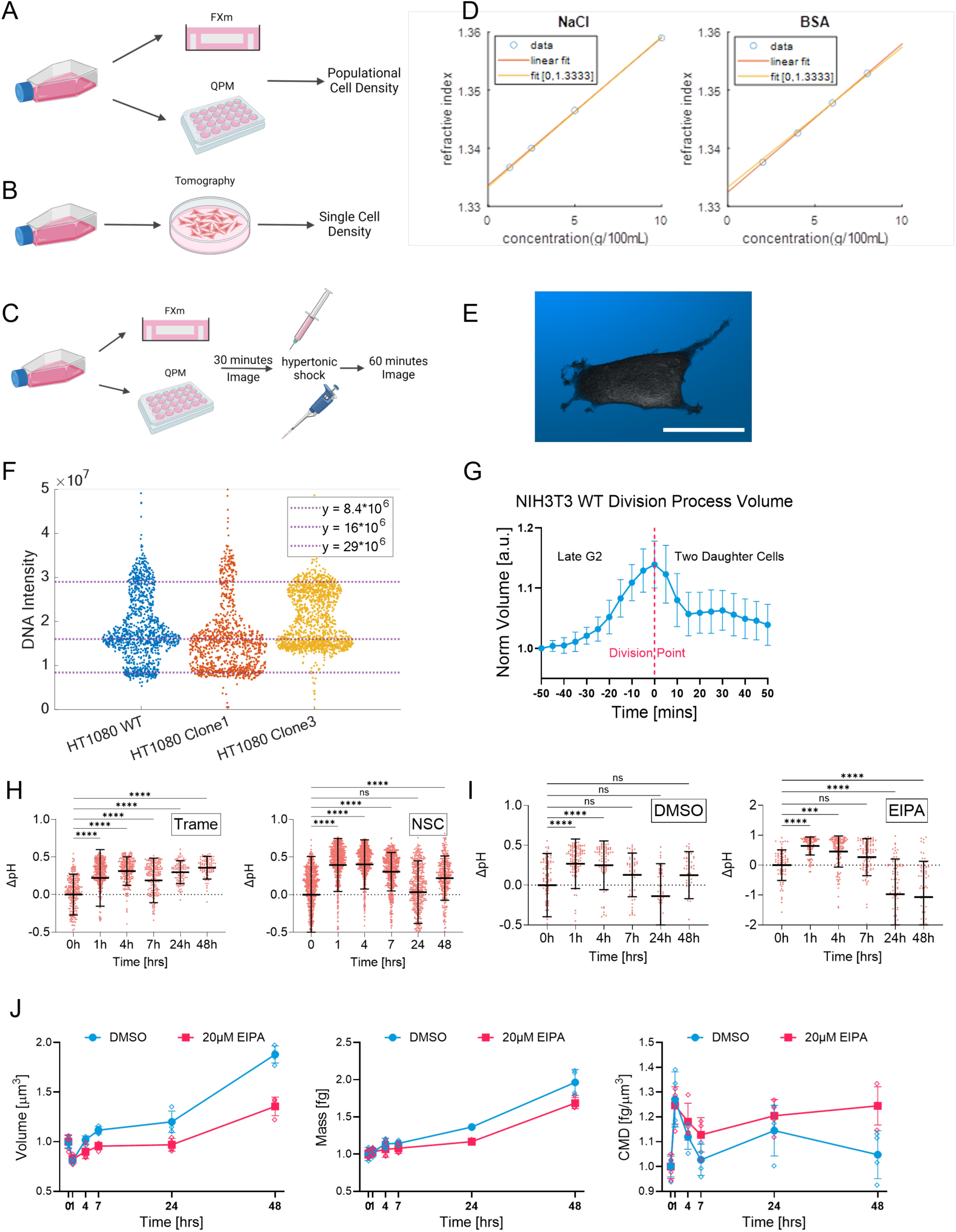
(A) A schematic of population cell volume, mass, and density experiment. (B) A schematic of holotomography single cell volume, mass, and density experiment. (C) A schematic of populational cell hyperosmotic shock volume, mass, and density experiment. (D) Refractive index versus mass density for NaCl and bovine serum albumin (BSA) water solutions. Each blue dot represents a single data point. N = 2. Yellow and red trendlines are linear fittings to the data points, which have a slope that represents the correlation constant between refractive index and solute mass density. (E) A representative holotomography 3D cell shape reconstruction. (F) DNA content of different HT1080 single cell clones quantified by Hoechst staining. n ≥ 1054. (G) NIH3T3 cell volume during mitotic swelling. We picked cells right before they separated into daughter cells to quantify the cell division volume, mass, and density. n = 39. (H) Intracellular pH change over time in hypertonic conditions with 100 nM Trame and 10 μM NSC. n ≥ 95. (I) HT1080 intracellular pH over time in hypertonic condition with DMSO and 20 μM EIPA. (J) HT1080 volume, mass, and density over time in hypertonic conditions treated with DMSO and 20uM EIPA. n ≥ 49. N = 4. (G, H, I, J) Data are presented as the mean ± standard deviation. (H, I) One-way ANOVA tests followed by Dunn-Šidák’s multiple-comparison test were conducted between data sets. *p^ns^* > 0.05, *p*\* ≤ 0.05, *p*\*\* ≤ 0.01, *p*\*\*\* ≤ 0.01, *p*\*\*\*\* ≤ 0.0001. (E) Scale bars, 30 μm.

**Supplemental Figure 2:**
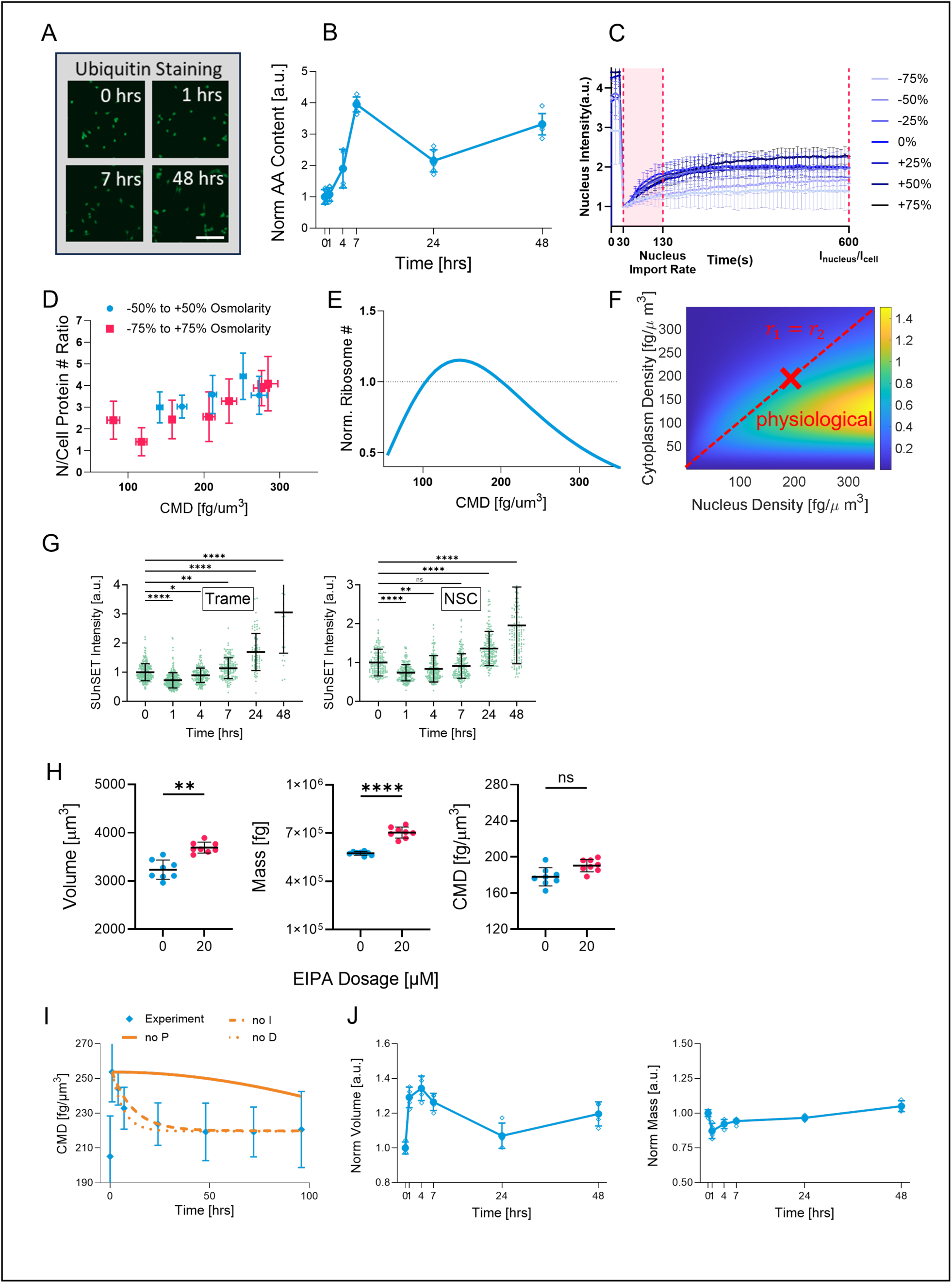
(A) Representative images of anti-ubiquitin staining in isotonic and 1, 7, 48 hrs in hypertonic condition. (B) Cell amino acid content over time in hypertonic condition. N = 6. (C) Normalized nucleus NLS-tdTomato- NES intensity FRAP trajectories with 2.5% FBS, and -75% to +75% osmolarity range. n ≥ 9. (D) Nucleus to cell NLS-tdTomato-NES content ratio 10 mins after the nucleus NLS-tdTomato-NES photobleaching. n ≥ 9. (E) Computed mature cytoplasmic ribosome numbers at different CMDs. (F) Heatmap showing theoretically simulated cell growth rate as a function of nucleus and cytoplasm mass density. Physiological densities are labeled as a red cross. The red dotted line shows the cytoplasmic density equal to the nucleus density. (G) Quantified SUnSET intensities over time in hypertonic conditions treated with 100nM Trame and 20µM NSC. n ≥ 29. (H) HT1080 volume, mass, and density in an isotonic environment treated with DMSO and 20 µM EIPA for 48 hrs. N = 8. (G) PID control model when deleting only one of the proportional, integral, and differential terms. (H) Cell volume and mass over time in 50% hypotonic condition. (B, C, D, G, H, I, J) Data are presented as the mean ± standard deviation. (G, H) One­way ANOVA tests followed by Dunn-Šidák’s multiple-comparison test were conducted between data sets. *p^ns^* > 0.05, *p*\* ≤ 0.05, *p*\*\* ≤ 0.01, *p*\*\*\* ≤ 0.01, *p*\*\*\*\* ≤ 0.0001. (A) Scale bars, 200 μm.

**Supplemental Figure 3:**
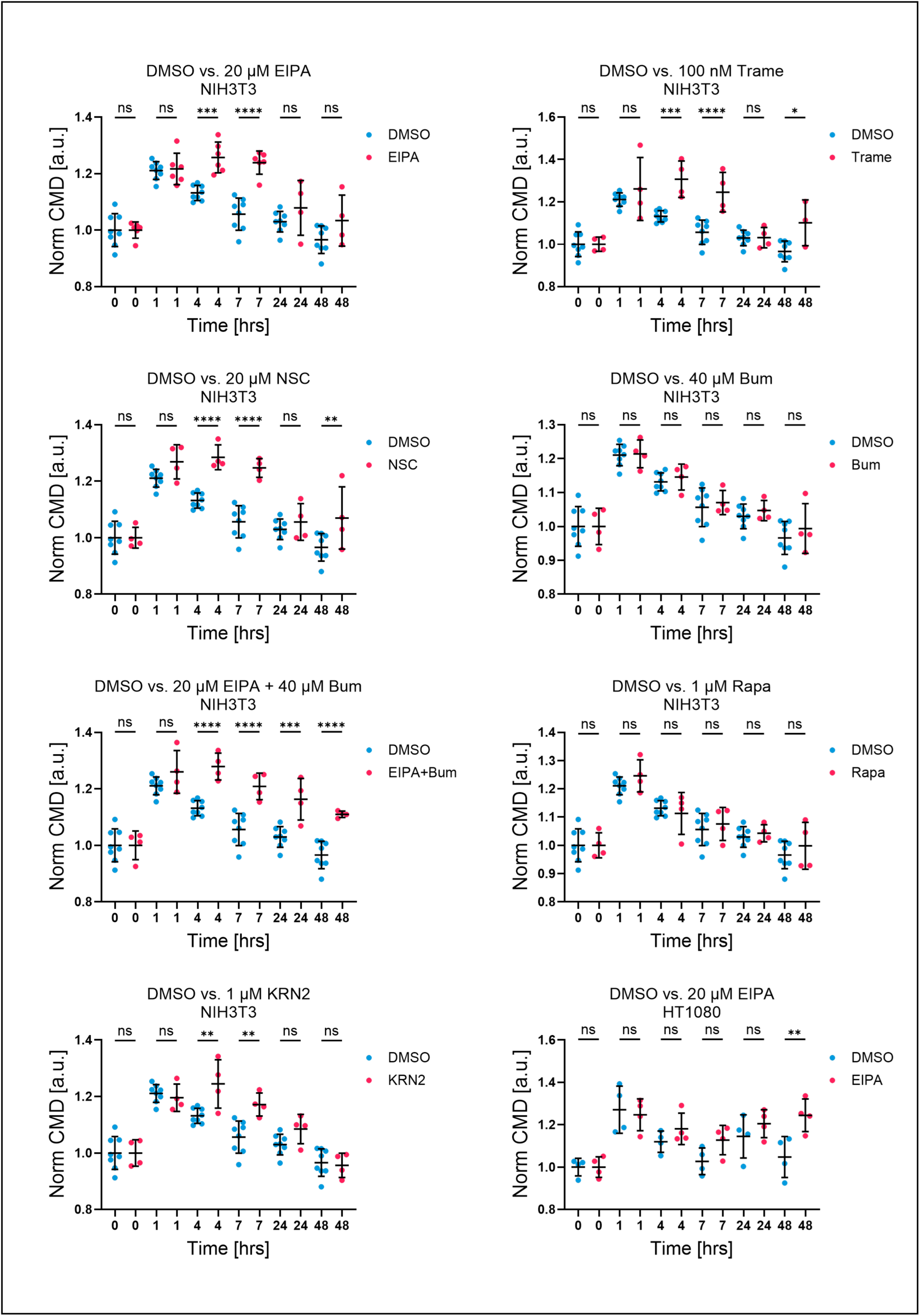
Statistical test for all NIH3T3 and HT1080 density over time under hypertonic stress experiments. Data are presented as the mean ± standard deviation. One-way ANOVA tests followed by Dunn-Šidák’s multiple-comparison test were conducted between data sets. *p^nS^* > 0.05, *p*\* ≤ 0.05, *p*\*\* ≤ 0.01, *p*\*\*\* ≤ 0.01, *p*\*\*\*\* ≤ 0.0001.

**Table s1:**
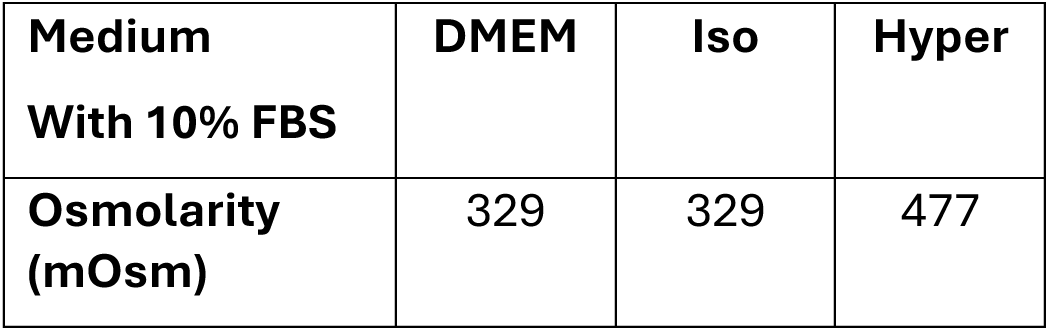
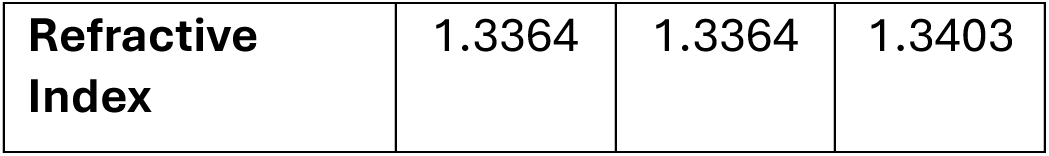
Cell Culture and Hypertonic Medium Osmolarity and Refractive Index.

**Table s2:**
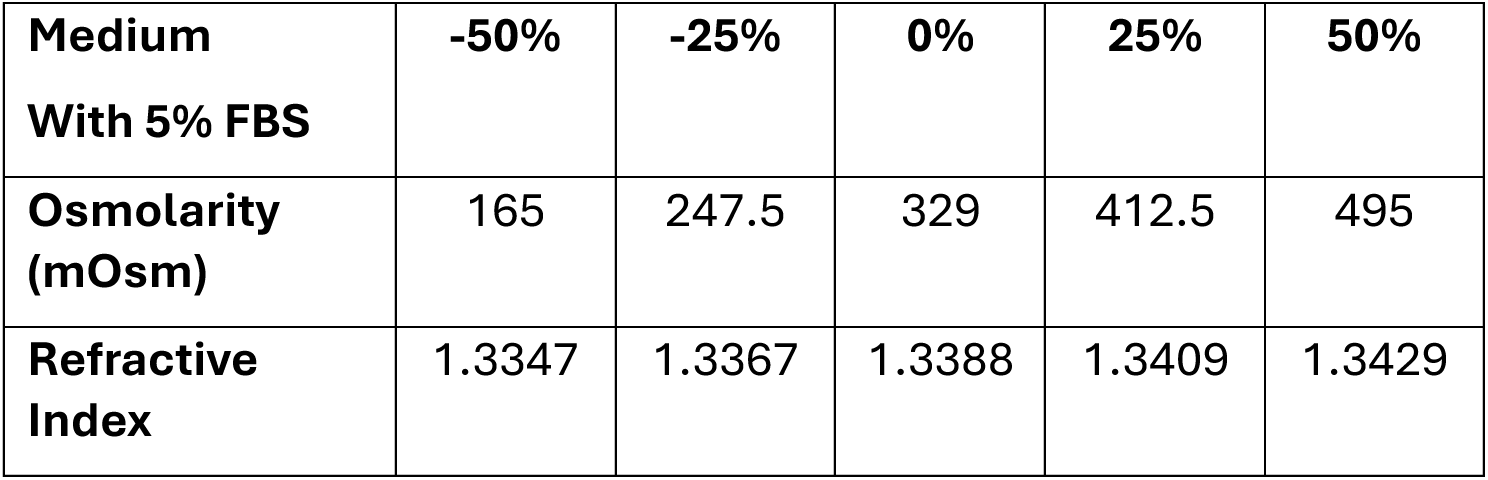
5% FBS Medium Osmolarity and Refractive Index.

**Table s3:** 2.5% FBS Medium Osmolarity and Refractive Index.

|  |  |  |  |  |  |  |  |
| --- | --- | --- | --- | --- | --- | --- | --- |
| **Medium With 2.5% FBS** | **-75%** | **-50%** | **-25%** | **0%** | **25%** | **50%** | **75%** |
| **Osmolarity (mOsm)** | 83 | 165 | 248 | 330 | 412 | 495 | 578 |
| **Refractive Index** | 1.3338 | 1.3359 | 1.3380 | 1.3400 | 1.3421 | 1.3441 | 1.3461 |

